# Spatiotemporal components of sustained functional hyperemia are differentially modulated by locomotion and silenced with vascular chemogenetics

**DOI:** 10.1101/2021.09.15.460557

**Authors:** Govind Peringod, Linhui Yu, Kartikeya Murari, Grant R Gordon

## Abstract

Neural activity underlying sensation, movement or cognition drives regional blood flow enhancement – termed functional hyperemia – to increase the oxygen supply to respiring cells for as long as needed to meet energy demands. However, functional hyperemia is often studied under anesthesia which typically yields response profiles that appear temporally and spatially homogenous. We have insufficient understanding of the underlying kinetics of oxygen delivery in awake animals, especially during specific behaviours that may influence neurally-driven enhancements in cerebral blood flow. Using widefield intrinsic optical signal imaging in awake, head-fixed but active mice, we demonstrated distinct early and late components to changes in intravascular oxygenation in response to sustained (30s) whisker stimulation. We found that the late component (20-30s), but not the early component (1-5s), was strongly influenced by level of whisking/locomotion in the region of highest response and in surrounding regions. Optical flow analyses revealed complex yet stereotyped spatial properties of the early and late components that were related to location within the optical window and the initial state of the cerebral vasculature. In attempt to control these complex response characteristics, we drove a canonical microvasculature constriction pathway using mural cell Gq-chemogenetic mice. A low-dose of systemic C21 strongly limited both the magnitude and spatial extent of the sensory-evoked hemodynamic response, showing that functional hyperemia can be severely limited by direct mural cell activation. These data provide new insights into the cerebral microcirculation in the awake state and may have implications for interpreting functional imaging data.

## INTRODUCTION

The brain’s microcirculation requires unique spatiotemporal regulation to meet the high energy demands of active neural circuits. At baseline, ongoing, high-frequency neuronal rhythms entrain low-frequency oscillations in contractile microvasculature (Mateo et al. 2017), which is thought to be important for the perfusion of blood to constantly and rhythmically active tissue (Cole 2019). Additional regulation is required to support spatially heterogenous changes in neural activity that drive the animal’s sensory, motor or cognitive response to external and internal cues (Iadecola 2017), termed ‘functional hyperemia.’ Most studies have examined cerebral hemodynamics under the context of functional hyperemia in anesthetized animals in response to sensory stimulation. When measured at a point location in this state, this limits the degree, kinetics and spatial aspects of the observed blood flow response (Gao et al. 2017, Tran and Gordon 2015b), and can provide misleading measurements of oxygenation that do not reflect the realistic supply dynamics in the awake brain (Lyons et al. 2016). Furthermore, in many previous studies that have examined the hemodynamic response to prolonged sensory stimulation (>10 s) in anesthetized/sedated animals, the kinetics of functional hyperemia were either not reported, or showed a square-like on-off response. In work examining neurovascular coupling in awake animals, the data indicate there are at least two components to the response (Sharp et al. 2015, Drew et al. 2011, Takuwa et al. 2012, Takuwa et al. 2013), with an initial phase (∼1-5 s) showing distinct kinetics compared to a later phase (>5 s), yet we know little about these putative phases or how they are regulated. As well, locomotion and locomotion-associated whisking are important behavioral states that drive neuromodulation of specific brain circuits (Paukert et al. 2014, Polack et al. 2013, Ding et al. 2013), Ayaz et al., 2019), elicit spatially heterogeneous increases in blood flow and tissue oxygenation (Huo et al. 2014, Zhang et al. 2019), and potentiate astrocytic Ca^2+^ signals associated with neurovascular coupling (Tran et al. 2018). Yet, the literature concerning distinct spatial and temporal contributions to functional hyperemia is limited (Cauli and Hamel 2010), especially in awake animals where there could be behavioural influences.

Understanding putative early and late components of functional hyperemia could be important because different experimental preparations (isolated vessel, brain slice, anesthetized and awake *in vivo*) have implicated several cell types and many cellular pathways in neurovascular coupling, with ‘redundancy’ often hailed as an explanation for the emergent complexity of this essential phenomenon. However, an alternative hypothesis is that distinct cell-types and pathways are recruited on specific timescales, at specific locations within the cerebrovascular tree, or within certain physiological contexts in the awake animal, to properly coordinate cerebral blood flow and oxygenation.

Here we used an awake mouse preparation adapted for widefield intrinsic optical signal (WF-IOS) imaging to explore the temporal and spatial properties of functional hyperemia under relatively realistic conditions *in vivo* (Tran and Gordon 2015a). We characterized changes to intravascular oxygenation in response to prolonged whisker stimulation in active mice and tested the hypothesis that spatial and temporal components of functional hyperemia are differentially regulated by the behavioral state and can be constrained by direct manipulation of the microvasculature.

## RESULTS

### Sustained sensory stimulation evokes two-component oxygenation changes

We monitored changes in intravascular oxygenation within the right barrel cortex in response to gentle, 30-second air-puff stimulation of the left vibrissae by measuring diffuse-reflectance changes of green (530 nm) and red (660 nm) wavelengths, and then estimating concentration changes in oxygenated (HbO) and deoxygenated (HbR) hemoglobin using standard methods (Ma et al. 2016). Since the 2D image formed at the camera is a superficially weighted sum of signals originating from the cortex, dura was removed during craniotomy to avoid contributions from extracortical vessels, which have previously been reported to constrict during locomotion (Gao and Drew 2016). We used mural cell (PDGFRβ-Cre x RCL-GCaMP6s) reporter mice but no Ca^2+^ data was acquired in this WF-IOS study. In leu of neural activity, we measured its ultimate output – behaviour. Animals were head fixation on an air-supported Styrofoam ball (Tran and Gordon, 2015) and permitted to run at will. General behaviour and whisking were captured with a dedicated camera, whereas locomotion activity was quantified by additionally videoing a region on the ball surface (Figure 1A-C; also see Methods). Mice were first trained for head fixation for several days before an acute cranial window was installed followed by immediate imaging once fully awake (Tran and Gordon, 2015) (see methods).

**Figure 1:**
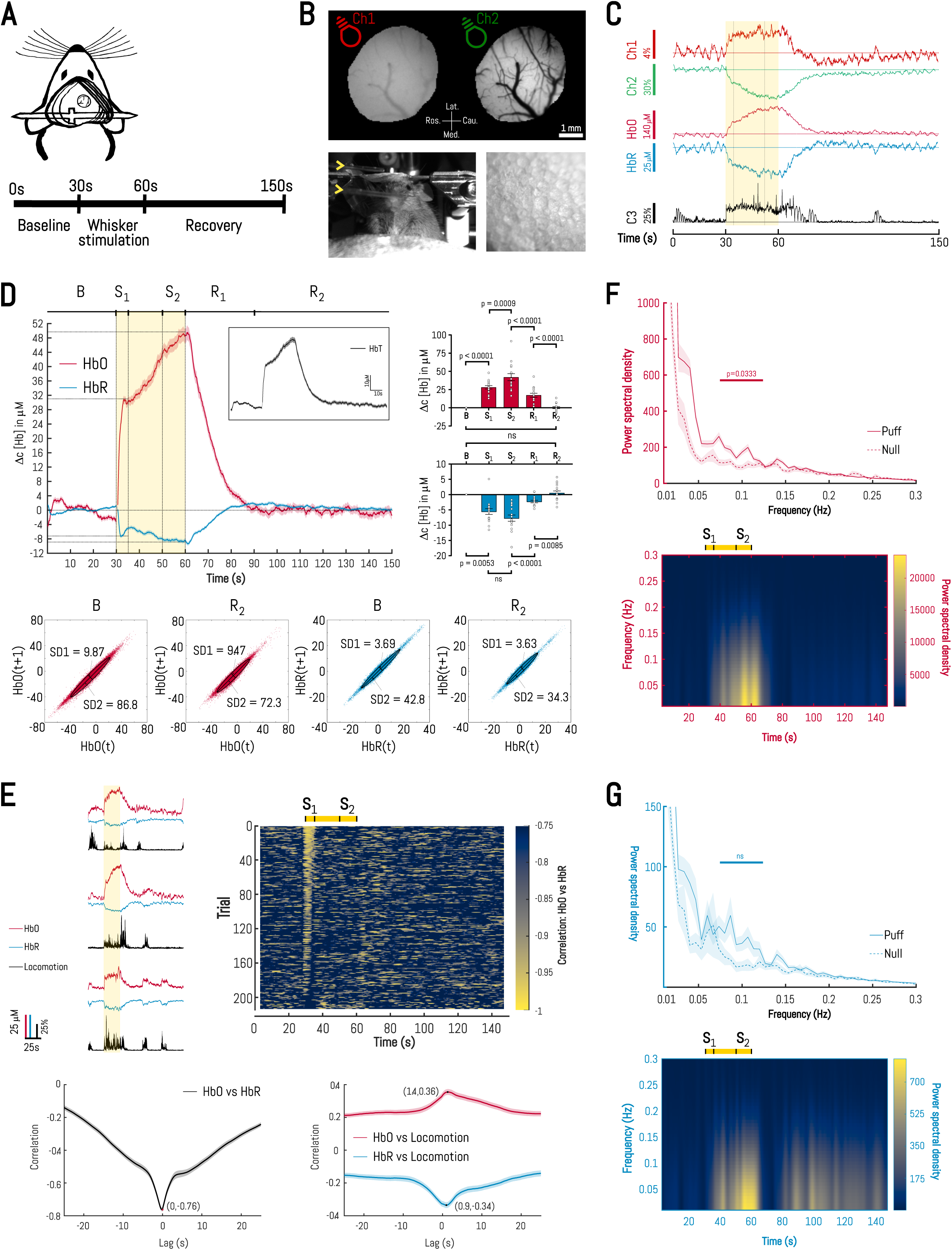
Sustained sensory stimulation evokes two-component oxygenation changes. A. Schematic of experimental setup: (Above) Animal preparation with headbar and cranial window; (Below) Imaging protocol. B. (Above) Representative images showing raw grayscale reflectance of cranial window under 660nm and 530nm illumination; (Below) Frames from behavioral cameras during a trial. Yellow arrowheads indicate source of air puff. ‘Ch1’: red channel; ‘Ch2’: green channel. Lat.: lateral; Med.: medial; Ros.: rostral; Cau.: caudal. C. Representative traces showing raw reflectance, hemodynamic and behavioral changes during whisker stimulation. ‘C3’: ball-facing behavioral camera. Yellow shading indicates whisker stimulation period. D. Trial-averaged time-course of hemodynamic changes during whisker puff trials. Data shown represents mean ± SEM. Yellow shading indicates whisker stimulus period. B: Baseline (0-30 s); S_1_: Early stimulation (30-35 s); S_2_: Late stimulation (50-60 s); R_1_: Early recovery (60-90 s); R_2_: Late recovery (90-150 s) (Inset) Scatter dot plots showing distribution of animal means (N=15 mice) for HbO and HbR changes in each temporal segment of the response. Error bars: mean change ± SEM. Repeated-measures one-way ANOVA with the Greenhouse-Geisser correction for HbO (F(1.346,20.2) =73.74, P<0.0001), Sidak’s multiple comparisons test: B vs S_1_ (P<0.0001, 95%CI -30.27 to -19.25); S_1_ vs S_2_ (P=0.0009, 95%CI -34.88 to -8.615); S_2_ vs R_1_ (P<0.0001, 95%CI 20.19 to 38.22); R_1_ vs R_2_ (P<0.0001, 95%CI 10.57 to 23.69); B vs R_2_ (P>0.9999, 95%CI -4.30 to 3.94). Repeated-measures one-way ANOVA with the Greenhouse-Geisser correction for HbR (F(1.394,20.92)=40.72, P<0.0001), Sidak’s multiple comparisons test: S_2_ vs R_1_ (P<0.0001, 95%CI -8.56 to -3.87); R_1_ vs R_2_ (P=0.0085, 95%CI -4.95 to -0.66); B vs R_2_ (P>0.9999, 95%CI -2.45 to 1.39). Friedman test for HbR (χ^2^(2)=24.88, P<0.0001) with Dunn’s multiple comparisons test: B vs S_1_ (P=0.0053); S_1_ vs S_2_ (P=0.1037). ‘ns’: not significant. E. (Above, left) Time-courses of hemodynamic activity and locomotion during individual whisker puff trials from three different animals; (Above, right) Heatmap of moving correlation between HbO and HbR time-courses for whisker puff trials. Colormap encodes correlation coefficients in the range [-1, -0.75] and yellow bar above indicates whisker stimulus period. (Below, left) Summary cross-correlation between HbO and HbR for whisker puff trials. (Below, right) Summary cross-correlation between hemodynamics and locomotion for whisker puff trials. Data shown is mean ± SEM. F. (Above) Trial-averaged single-sided amplitude spectra of HbO changes during null and whisker puff trials; (Below) Trial-averaged spectrogram of HbO activity during whisker puff trials. Yellow bar above indicates whisker stimulus period. Paired two-tailed t-test, t(14)=2.36, P=0.0333. G. (Above) Trial-averaged single-sided amplitude spectra of HbR changes during null and whisker puff trials; (Below) Trial-averaged spectrogram of HbR activity during whisker puff trials. Yellow bar above indicates whisker stimulus period. Paired two-tailed t-test, t(14)=0.08, P=0.9343. ‘ns’: not significant.

HbO and HbR responses to sustained whisker stimulation (214 trials, 16 mice) displayed two putative components with an initial rapid rise to a first peak (S_1_), followed by a slower increase to a second, higher peak near the end of the stimulation (S_2_); this two-peak profile was also seen in the total blood volume (HbT) curves (Figure 1D). The ratio of the peaks S_2_/S_1_ was greater for HbO (1.91±0.16) than HbR (1.27±0.25), perhaps indicating enhanced vasodilation and oxygen delivery for the late component of functional hyperemia. Both HbO and HbR fully returned to baseline values over a decay period (R_1_) that was slower than the rising periods during stimulation (S_1_ and S_2_) (HbO: τ_S1_ = 2.42±0.31 s; τ_S2_ = 3.7±1.28 s; τ_R1_ = 10.55±1.05 s. HbR: τ_S1_ = 1.83±0.26 s; τ_S2_ = 3.24±1.25 s; τ_R1_ = 8.36±1.18 s). Notably, trial-to-trial variability was significantly lower during the late recovery period (R_2_) than during the baseline period (B); this was evident in the ellipse-fitted Poincaré maps 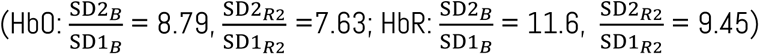, and in the comparison of the coefficients of variation for each time-series (median of differences: ΔHbO = -0.0348 µM, p = 0.0354; ΔHbR = 0.710 µM, p = 0.0175), suggesting a vessel refractory period after long stimulations (Cannestra et al. 1998).

In the awake mouse, these stimulus-evoked changes were superimposed upon a dynamic baseline consisting of rhythmic, low-amplitude fluctuations that repeated once every ∼10 seconds (Mayhew et al. 1996) related to vasomotion (Mateo et al. 2017), and one or more bouts of naturally occurring functional hyperemia during whisking/locomotion events (Tran et al. 2018) (Figure 1E). These latter events varied in length, were distributed randomly within the recordings, and consisted of a sustained deviation from baseline. Correlation analyses showed that HbO and HbR activity were in phase and strongly anticorrelated (R < -0.75), and that both signals lagged locomotion activity by a time delay (HbO: 1.4 s; HbR: 0.9 s), which is consistent with functional hyperemia. Intriguingly, we observed spikes in the time-resolved correlation between HbO and HbR during S_1_ and immediately after the end of 30-second whisker stimulation (Figure 1E), which occurred due to concomitant increases in oxygen consumption and delivery during these time periods. Such changes may reflect neurons registering the onset and offset of the stimulus (Kyriazi et al. 1994).

Frequency domain analyses showed that most of the spectral power was concentrated at very low frequencies (0.01-0.3 Hz), and that 30 sec whisker stimulation elicited an expected increase in spectral power during the stimulation period in this frequency range (Figure 1F-G). Comparing whisker-stimulation trials with trials in which no whisker puff stimulus was given (‘null trials’) revealed that the spectral power within an ultra-low-frequency band (0.075-0.125 Hz) increased significantly for HbO (p = 0.0333) during whisker stimulation, but not for HbR (p = 0.9343). A similar divergence, also specific to HbO (p = 0.0463), was observed in a higher frequency window (1.75-2.25 Hz) that is likely related to the animal’s inspiration-expiration cycle (Zhang et al. 2019) (Supplementary Figure 1).

### Oxygenation changes exhibit a stereotyped spatial pattern

Visual inspection of HbO and HbR time-stacks from whisker-puff trials showed changes in signal intensity originating in a localized active region and propagating to surrounding vascular structures within ∼2 seconds of stimulus onset (see Movie 1). Typically, there was only one active region within a given 3×3 mm optical window, and the distinction between active and surrounding regions was more apparent in S_2_ than in S_1_, and in HbO than in HbR time-stacks, likely because both S_2_ and HbO exhibit comparatively larger magnitude changes (Figure 2A). Despite variability from trial to trial, whisker stimulation and locomotion-associated hemodynamics both routinely elicited a visible dilation of pial arteries and caused associated increases in [HbO] within pial arteries and decreases in [HbR] within pial veins. These changes tracked each other and were apparent for ∼5 seconds after stimulus offset before slowly returning to baseline over ∼25 seconds.

**Figure 2:**
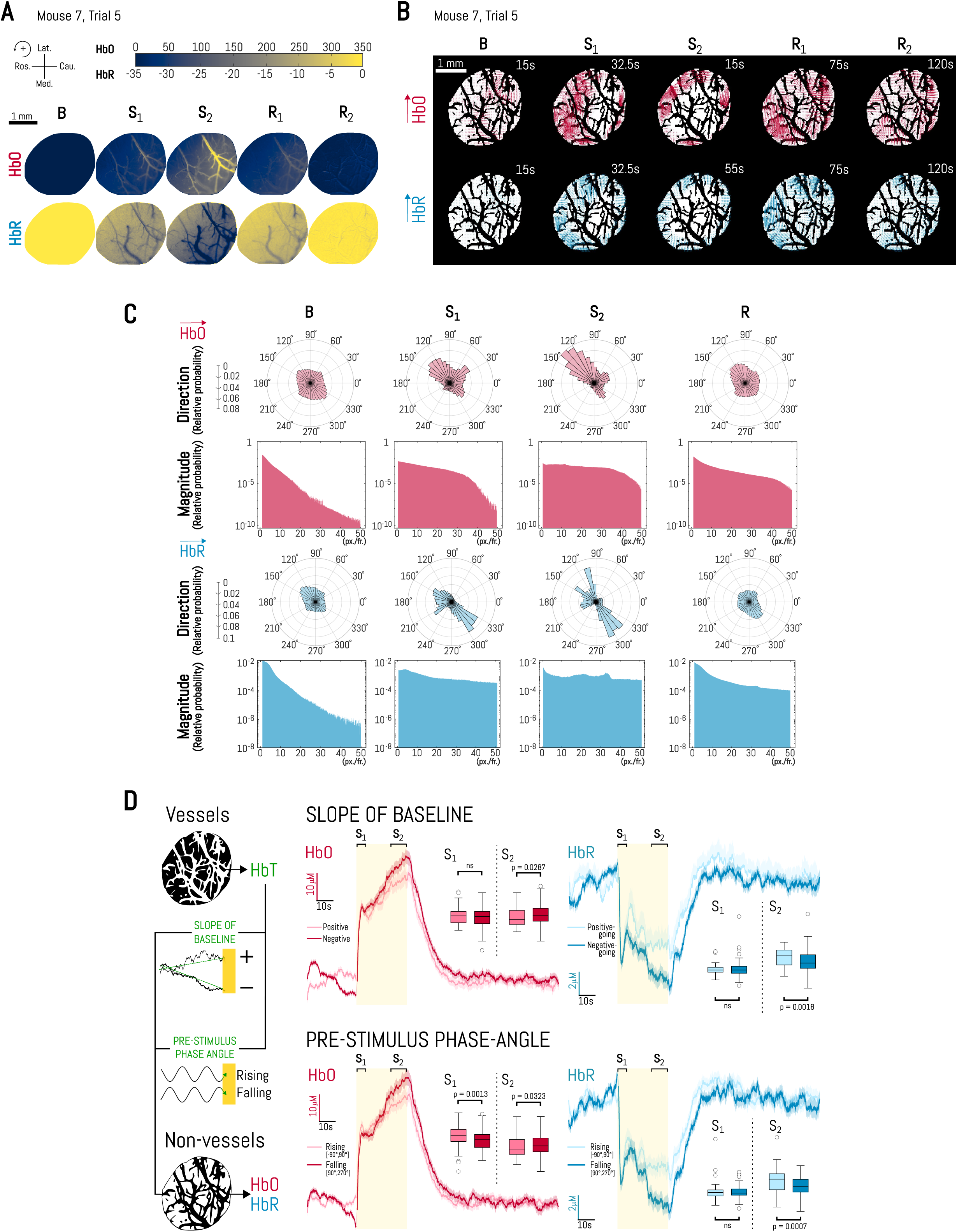
Oxygenation changes exhibit a stereotyped spatial pattern. A. Mean intensity projections from five different segments of the stimulus-evoked hemodynamic response for a representative whisker puff trial. Colormap encodes pixel-by-pixel Δc[Hb] values in μM and is limited to the range [0, 350] for HbO and [-35, 0] for HbR. Axis for spatial measurements is shown (top left). B: Baseline (0-30 s); S_1_: Early stimulation (30-35 s); S_2_: Late stimulation (50-60 s); R_1_: Early recovery (60-90 s); R_2_: Late recovery (90-150 s) B. Representative frames from pixel-by-pixel optical flow analysis for the same whisker puff trial as in 2A. Size of arrows denotes flow vector magnitude, and origin of arrows indicate pixel centers. C. Standard and polar histograms of the full dataset summarizing, respectively, magnitude and direction of optical flow vectors for 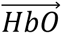 and 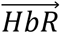 from trial-averaged time-stacks. D. (Above) Slope of baseline (positive vs negative) and pre-stimulus phase angle (-90° to 90° vs 90° to 270°) of the vessel-associated HbT trace was used to segregate non-vessel-associated HbO and HbR traces. In the representative vessel and non-vessel binary masks, pixels used for generating hemodynamic traces are shown in white. (Middle) Trial-averaged time-courses of hemodynamic changes for whisker stimulation trials with a positive-going or negative-going baseline HbT trace. Data shown represents mean ± SEM. Yellow shading shows whisker stimulus period. (Inset) Tukey box-and-whiskers-plots of trial means (from 14 mice) for HbO and HbR changes during S_1_ and S_2_. Mann-Whitney two-tailed unpaired tests, HbO S_1_: U=3757; P=0.4065; HbO S_2_: U=3279; P=0.0287; HbR S_1_: U=3889, P=0.6475. HbR S_2_: U=2952, P=0.0018. ‘ns’: not significant (Below) Trial-averaged time-courses of hemodynamic changes for whisker stimulation trials with a rising or falling phase in the HbT trace immediately before the stimulus period. Data shown represents mean ± SEM. Yellow shading shows whisker stimulus period. (Inset) Tukey box-and-whiskers-plots of trial means (from 14 mice) for HbO and HbR changes during S_1_ and S_2_. Error bars: mean change ± SEM. Mann-Whitney two-tailed unpaired tests, HbO S_1_: U=3069; P=0.0013; HbO S_2_: U=3455; P=0.0323; HbR S_1_: U=4193, P=0.9252. Welch-corrected two-tailed unpaired t-test, HbR S_2_: t(181)=3.445, P=0.007. ‘ns’: not significant.

To describe the observed two-dimensional displacements of surface brightness patterns more quantitatively, we performed dense optical flow analysis on HbO and HbR time-stacks to compute instantaneous velocities at every pixel within the optical window (Figure 2B; also see Movies 2-3). Visual inspection confirmed that changes in HbO and HbR velocity vector fields 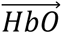 and 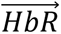 respectively) coincided in space and time with observed brightness changes in corresponding time-stacks. Both fields were characterized by smooth, circular flow patterns with streamlines that largely adhered to the curvilinear boundaries of surface vessels. Magnitudes of 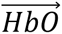 vectors were consistently larger than 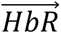 vectors and, intriguingly, 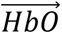 vectors were almost always rotated by 180° relative to 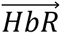 vectors (i.e. brightness displacements in HbO and HbR time-stacks were directionally opposed to each other). Summary histograms of the entire dataset (214 trials, 16 mice) showed a stereotyped response (Figure 2C): velocity vectors were small and randomly oriented during the baseline period (B: 0-30 s) but the stimulation period (S: 30-60 s) saw a dramatic increase in vector magnitudes and their acquisition of a directional bias with respect to the rostral-caudal axis (HbO: +135°, HbR: –45°); the recovery period (R: 60-150 s) was associated with a return to baseline profile of vector directions and (to a lesser extent) vector magnitudes. Within the stimulation period, vectors in S_2_ had higher vector magnitudes and a more pronounced directional bias. These data show a stereotyped directionality of oxygenation changes which likely reflect vascular topology of oxygenated blood entering solely from the MCA laterally and deoxygenated blood draining towards the brain midline via more numerous vein collection routes (Hirsch et al. 2012, Xiong et al. 2017).

We then asked whether the initial state of the vessel contributed to the trial-to-trial variability observed in the two-component oxygenation response to 30-second whisker stimulation (Figure 2D). We visually identified locations of pial arteries within each cranial window and grouped individual trials according to the nature of the total hemoglobin response (HbT = HbO + HbR) within this ‘vessel’ region before stimulus onset (184 trials, 14 mice). Specifically, we calculated: 1) the slope of the HbT response during the baseline period (0-30 s), which was classified as being either positive or negative; and 2) the phase-angle of oscillation in the half-second immediately preceding the whisker puff stimulus, which was classified as being either rising (-90° to 90°) or falling (90° to 270°) (Saka et al. 2012). We used these grouping variables to compare how the initial state of pial arteries influenced HbO and HbR responses in non-pial vessel regions during S_1_ and S_2_. A negative baseline HbT slope and a falling HbT phase-angle produced a larger parenchymal HbR response during S_2_ (Hodges-Lehmann difference between medians of slope groups: 6.96 µM, p = 0.0018; difference between phase-angle groups: 2.98±0.87 µM, p = 0.0007), but not during S_1_. We observed the same trend for HbO with respect to the slope of HbT baseline; however, a falling HbT phase-angle produced a smaller HbO S_1_ response but a larger HbO S_2_ response as compared to a rising HbT phase-angle (Hodges-Lehmann difference between medians, S_1_: 4.11 µM, p = 0.0013; S_2_: 6.43 µM, p = 0.0323). These data suggest that a constricting vessel at the onset of whisker stimulation largely enhances the magnitude of hemodynamic change during the late component of functional hyperemia (S_2_), which is more pronounced for HbR than HbO.

Taken together, these data show a stereotyped spatial pattern of two-component blood flow and tissue oxygenation responses in the barrel cortex, with variations in response magnitude that are related to the initial conditions of the vasculature and location within the cranial window.

### Late component of functional hyperemia is influenced by whisking/locomotor state

Behavioral states like arousal and locomotion have widespread modulatory effects on the activity of cells in the cortex and are associated with regional increases in cerebral blood volume and oxygenation (Huo et al. 2014, Paukert et al. 2014, Polack et al. 2013). We first asked whether the initial changes in S_1_ were different in puff-induced versus voluntary locomotion bouts of hemodynamic activity (Figure 3A). We computed locomotion-triggered averages by aligning intrinsic optical signals to the onset of locomotion using only running events ≥ 10 s in duration that were preceded by a stable 10 s baseline (28 puff-induced bouts across 9 mice, 190 voluntary bouts across 21 mice). In both cases, HbO and HbR changed significantly from baseline (p < 0.0001) within ∼3 s after locomotion onset (Figure 3B); however, the magnitude of hemodynamic changes was significantly larger for puff-induced running as compared to voluntary running (p < 0.0020). As a potential explanation for this difference, we compared the rise-times of averaged locomotion traces and found that whisker puff-induced faster running than voluntary bouts (p = 0.0067).

**Figure 3:**
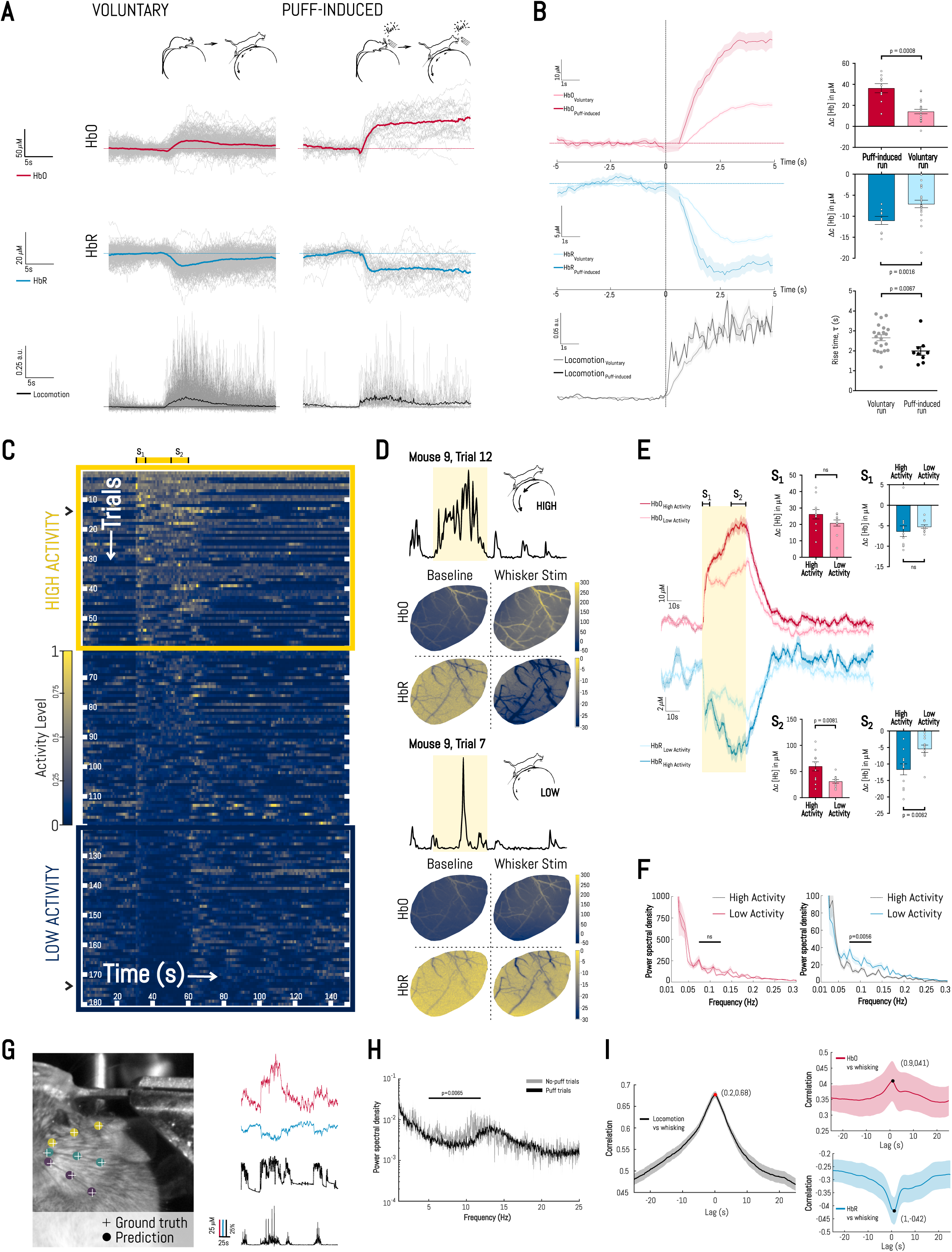
Late component of functional hyperemia is influenced by the whisking/locomotor state. A. Time-course of hemodynamic changes aligned to the onset of whisker puff stimulus for: (Left) Voluntary bouts of locomotion; (Right) Whisker puff-induced bouts of locomotion. Color trace is mean, gray traces are raw data (190 voluntary bouts, 28 whisker puff-induced bouts). B. (Left) Trial-averaged time-courses of hemodynamic changes during voluntary and whisker puff-induced bouts of locomotion, centered at the onset of the whisker puff stimulus. Data shown represents mean ± SEM. (Right) Scatter dot plots showing distribution of animal means for HbO and HbR changes for voluntary (N=21 mice) and whisker puff-induced (N=9 mice) locomotion bouts. Error bars: mean change ± SEM. Paired two-tailed t-tests, HbO: t(8)=5.239, P=0.0008; HbR: t(8)=4.681, P=0.0016. Mann-Whitney two-tailed unpaired test, Locomotion: U=36, P=0.0067. C. Heatmap of locomotion activity for a subset of whisker puff trials that was selected for activity-level comparisons. Traces are sorted in descending order of total activity during stimulation period (30-60 s). Colormap encodes relative activity in the range [0, 1]. Yellow boxes indicate trials designated as ‘high-activity’ and ‘low-activity’. Blue arrow markers identify trials selected as representative examples. Yellow bar above shows whisker stimulation period. D. Representative locomotion activity outputs from a ‘high-activity’ trial (top) and a ‘low-activity’ trial (bottom). For each: above is the trace of relative locomotion activity; below is the comparison of mean intensity projections of baseline (left) and stimulation (right) periods from trial-averaged HbO- and HbR timeseries. Colormap encodes pixel-by-pixel Δc[Hb] values in μM and is limited to the ranges [- 50, 300] for HbO and [-30, 0] for HbR. E. Trial-averaged time-courses of hemodynamic changes for low and high locomotion activity trials. Data shown represents mean ± SEM. Yellow shading shows whisker stimulus period. (Insets, right) Scatter dot plots showing distribution of animal means for changes during S_1_ (above) and during S_2_ (below) in high-activity (N=11 mice) and low-activity (N=11 mice) trial groups. Error bars: mean change ± SEM. Wilcoxon two-tailed matched-pairs signed rank tests, HbO S_1_: W=-32, P=0.1748; HbR S_1_: W=30, P=0.2061. Paired two-tailed t-tests, HbO S_2_: t(10)=3.29, P=0.0081; HbR S_2_: t(10)=3.454, P=0.0062. ‘ns’: not significant F. Trial-averaged single-sided amplitude spectra of hemodynamic changes in low and high locomotion activity trials. Mann-Whitney two-tailed unpaired tests, HbO: U = 28; P=0.2973; HbR: U=10, P=0.0056. ‘ns’: not significant G. (Left) Test image with labels indicating ground-truth and predicted pixel locations for three whiskers. (Right) Time-courses of hemodynamic activity and whisking from a whisker puff trial. H. (Above) Trial-averaged single-sided amplitude spectra of whisking activity for puff and no-puff trials. Mann-Whitney two-tailed unpaired test, U = 336; P=0.0065. I. Summary cross-correlation between (left) locomotion and whisking, (above, right) HbO activity and whisking, (below, right) HbR activity and whisking, for whisker puff trials. Data shown is mean ± SEM.

We then asked whether the level of locomotion activity affected the spatial and temporal profiles of the observed hemodynamic response during whisker-puff trials. We sorted a subset of the data (180 trials, 11 mice) by descending order of total activity within the 30-second whisker stimulation period (Figure 3C). Visual inspection of mean intensity projections of the optical window during the stimulation period showed that hemodynamic changes relative to baseline were consistently more pronounced for a given mouse in high locomotion activity trials as compared to low locomotion activity trials (Figure 3D). Major vessels were more visibly dilated, and the magnitude of HbO and HbR changes were larger across the optical window in high activity states. Strikingly, the magnitude of HbO and HbR changes in S_2_ was strongly influenced by animal locomotion (mean of differences: ΔHbO = 28.75±8.74 µM, p = 0.0081; ΔHbR = 6.25±1.81 µM, p = 0.0062), yet there was no significant difference between these measures during S_1_ (Figure 3E). Thus, increased locomotor activity preferentially increases CBF and oxygenation during the late component of functional hyperemia. Finally, frequency domain comparisons of these signals showed that the spectral content in the ultra-low-frequency band (0.075-0.125 Hz) was significantly attenuated by increased locomotion activity for HbR (p = 0.0056), but not for HbO (Figure 3F). This is consistent with higher oxygen consumption within the barrel cortex during high locomotion activity trials.

Since running is strongly associated with natural whisking in awake mice (Sofroniew et al. 2014), we also quantified single-whisker movements in three mice using a deep-learning-based, marker-less pose estimation software (Mathis et al. 2018, Nath et al. 2019, Mathis et al. 2019). Despite the poor and time-varying contrast between whiskers and background, DeepLabCut robustly identified two or three specified whiskers in each frame (Figure 3G). Frequency domain analysis showed that whisker-stimulation suppressed natural whisking frequencies in the 5-12 Hz (p = 0.0065) (Figure 3H). Cross-correlation analyses showed that locomotion lagged whisking by 0.2 seconds, and that whisking had a higher correlation with hemodynamics than locomotion but the same lag (∼1s) (Figure 3I).

### Center-surround pattern of hemodynamic response is modulated by locomotion activity

Whisker stimulation influences the balance of excitatory and inhibitory cellular activity in the neocortex, as well as a dynamic equilibrium between constrictory and dilatory activity of regional vessels (Kleinfeld et al. 2011). Several studies have demonstrated that this establishes an antagonistic center-surround spatial pattern of neural and hemodynamic activity in anesthetized and awake animals, with the ‘center’ region showing net neuronal depolarization and increased oxygenation (Devor et al. 2008, Devor et al. 2007, Devor et al. 2005, Knutsen et al. 2016, Uhlirova et al. 2016). Yet, it remains unclear whether this pattern persists during prolonged stimulation and whether it is affected by the level of locomotion activity within a given trial. We created ‘center’ and ‘surround’ masks for each animal by computing mean intensity image projections of HbO time-stacks for every trial and then identifying a region within the optical window that was consistently activated across all trials using histogram-based thresholding (Figure 4A). Since the ‘surround’ region was on average ∼4x larger than the ‘center’ region, we normalized HbO and HbR activity by area fraction (A_Region_/A_Total_) for comparisons between regions. Although the magnitude of the response was significantly higher for HbO and HbR in the center region during S_1_ and S_2_ as compared to the surround (p < 0.0100), the kinetics of the response as well as the S_2_:S_1_ magnitude ratio were similar in center and surround regions (Figure 4B). Interestingly, two-way ANOVAs revealed that there was significant interaction between the hemodynamic measure (HbO, HbR) and location within the window (center, surround), which was more pronounced in S_1_ (p < 0.0001) than in S_2_ (p = 0.0010). Thus, the center-surround effect was stronger for HbO than for HbR particularly during S_1_, which was also readily apparent in the summary traces.

**Figure 4:**
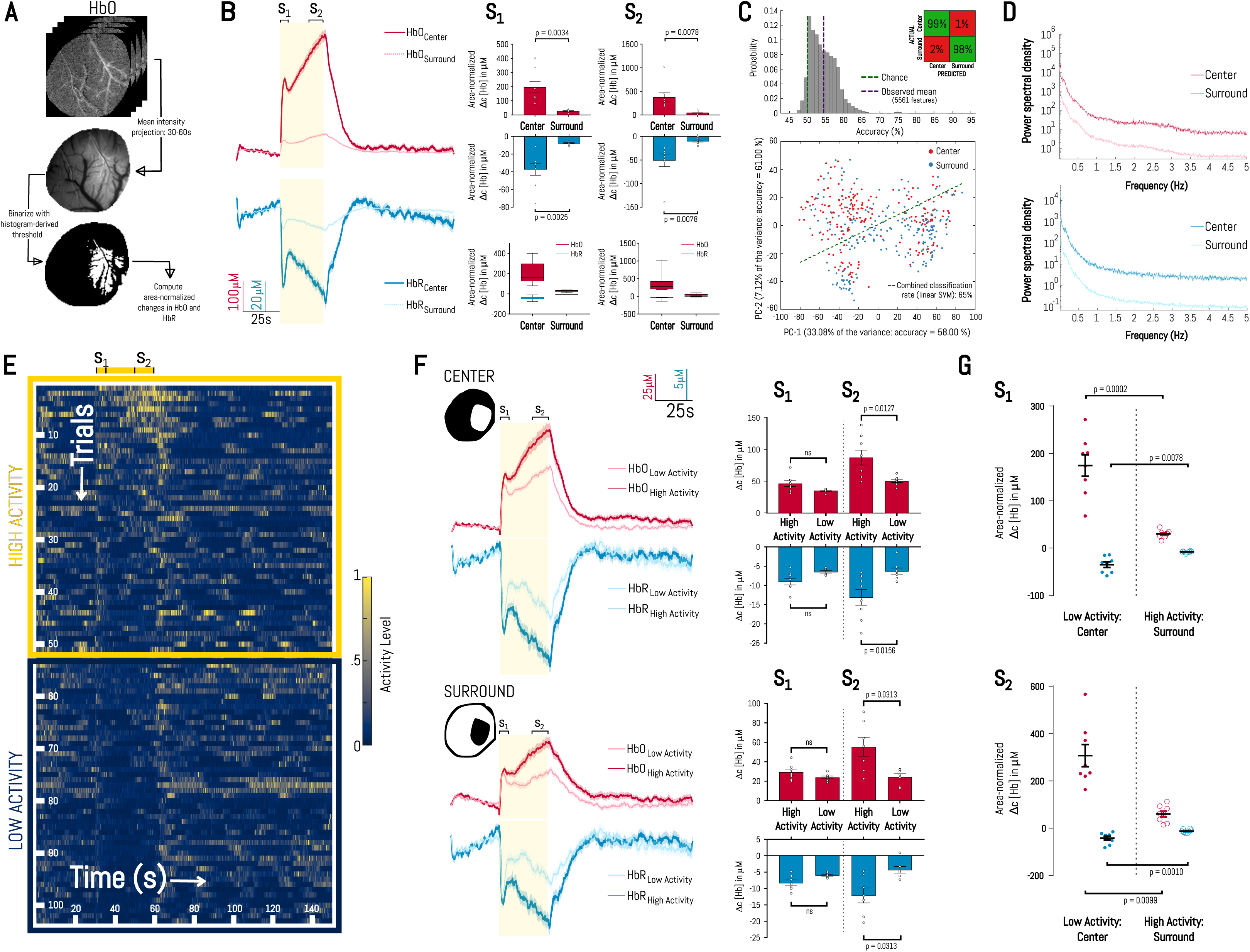
Center-surround pattern of hemodynamic response is modulated by locomotion activity. A. Schematic showing how frames from whisker stimulus period of HbO times-stacks were flattened and thresholded to create center-surround masks for each mouse. HbO and HbR changes were normalized by area fraction (A_region_/A_total_) for all comparisons between regions. B. (Left) Trial-averaged time-courses of hemodynamic changes in center and surround regions from whisker puff trials. Data shown represents mean ± SEM. Yellow shading shows whisker stimulus period. (Right, above) Scatter dot plots showing distribution of animal means for HbO and HbR changes in center (N=8 mice) and surround (N=8 mice) regions during S_1_ and S_2_. Error bars: mean change ± SEM. Paired two-tailed t-tests, HbO S_1_: t(7)=4.33, P=0.0034; HbR S_1_: t(7)=4.59, P=0.0025. Wilcoxon two-tailed matched-pairs signed rank tests, HbO S_2_: W=-36, P=0.0078; HbR S_2_: W=36, P=0.0078. (Right, below) Results of 2-way ANOVA between hemodynamic measure (HbO, HbR) and location within optical window (Center, Surround). S_1_: Interaction was significant (P<0.0001), accounting for 22.1% of the observed variance. S_2_: Interaction was significant (P=0.0010), accounting for 18.5% of the observed variance. C. (Above) Distribution of classification accuracies across 5561 features; (inset) confusion matrix for the linear SVM classifier used to separate ‘center’ and ‘surround’ hemodynamic time-courses. (Below) Low-dimensional representation showing the first two principal components of the classifier. D. Trial-averaged single-sided amplitude spectra in center and surround regions, for HbO changes (above) and HbR changes (below). E. Heatmap of locomotion activity for subsets of whisker puff trials that were selected for locomotion activity-level comparisons. Traces are sorted in descending order of total activity during stimulation period (30-60 s). Colormap encodes relative activity in the range [0, 1]. Yellow and blue boxes indicate trials designated as ‘high-activity’ and ‘low-activity’, respectively. Yellow bar above shows whisker stimulation period. F. (Left) Trial-averaged time-courses of hemodynamic changes in center and surround regions for low and high locomotion activity whisker puff trials. HbO and HbR changes were <u>not</u> normalized by area fraction for these within-region comparisons. Data shown represents mean ± SEM. Yellow shading shows whisker stimulus period. (Right) Scatter dot plots showing distribution of animal means for HbO and HbR changes in center (N=8 mice) and surround (N=7 mice) regions. Error bars: mean change ± SEM. Wilcoxon two-tailed matched-pairs signed rank tests, Center HbO S_1_: W=-28, P=0.0547; Center HbR S_1_: W=28, P=0.0547; Center HbR S_2_: W=34, P=0.0156; Surround HbO S_1_: W=- 12, P=0.3750; Surround HbO S_2_: W=-26, P=0.0313; Surround HbR S_1_: W=18, P=0.1563. Surround HbR S_2_: W=26, P=0.0313. Paired two-tailed t-test, Center HbO S_2_: t(7)=3.32, P=0.0127. ‘ns’: not significant G. Scatter dot plots comparing distribution of animal means for HbO and HbR changes in the center region of ‘low activity’ trials to the ‘surround’ region of ‘high activity’ trials, for S_1_ (above) and S_2_ (below). Paired two-tailed t-tests, HbO S_1_: t(7)=6.95, P=0.0002; HbO S_2_: t(7)=5.45, P=0.0010; HbR S_2_: t(7)=3.51, P=0.0099. Wilcoxon two-tailed matched-pairs signed rank tests, Center HbR S_1_: W=36, P=0.0078.

We asked whether center and surround regions within the optical window had distinct hemodynamic signatures. We used a MATLAB-based massive feature extraction framework to acquire 5561 unique features from each hemodynamic trace, and subsequently trained a linear classifier to separate ‘center’ from ‘surround’ time-series with >99% accuracy (Figure 4C). The most discriminative features were spread-dependent measures of raw and kernel-smoothed distributions of the time-series data, followed by measures of correlation, entropy and spectral power. Interestingly, the highest-performing features in this discrimination task were statistical measures commonly used for analyzing variations in the beat-to-beat interval of heart rate (e.g. pNN50), and goodness-of-fit measures (e.g. normalized error) when fitting distributions or models to the data. Follow-up frequency domain analyses also showed that power spectral density was persistently higher in the center region for both HbO and HbR (Figure 4D).

We then repeated activity-level comparisons, this time within center and surround regions (Figure 4E-F). The magnitude of HbO and HbR changes was strongly influenced by animal locomotion in S_2_, but not S_1_, in both center and surround regions (p < 0.0500). However, hemodynamic changes in the center region during low-activity trials was significantly larger (p < 0.0100) than corresponding changes in the surround region during high locomotion activity trials in both S_1_ and S_2_ (Figure 4G). This suggests that, in the awake animal, center-surround antagonism dominates over the modulatory effect of locomotion to maintain the spatial profile of the hemodynamic response.

### A canonical vasoconstriction pathway constrains functional hyperemia spatially and temporally

We examined whether a direct vascular manipulation altered the spatial and temporal properties of the two-component blood flow and oxygenation response. We drove the canonical Gq-IP_3_-signaling pathway in mural cells (vascular smooth muscle cells and pericytes) using chemogenetics, to cause active and ongoing vasoconstriction. To do this, we generated transgenic mice expressing <u>Gq</u>-coupled <u>d</u>esigner receptors <u>e</u>xclusively <u>a</u>ctivated by <u>d</u>esigner <u>d</u>rugs (Gq-DREADDs) specifically in mural cells (Figure 5A) (Armbruster et al. 2007). Spontaneous behaviors and associated hemodynamic activity in naïve PDGFRβ- Cre x CAG-LSL-GqDREADD mice during training sessions were similar to those we observed in PDGFRβ × GCaMP6s^+/–^ mice. We activated Gq-IP3-signaling selectively in mural cells of these mice (n=3) via intraperitoneal injection of a low dose (0.1 mg/kg) of the potent and selective DREADD agonist Compound 21 (C21) (Jendryka et al. 2019, Chen et al. 2015), which elicited a visible constriction of pial arterioles (Supplementary Figure 2A). This was either from direct vasoconstriction, or a myogenic response from increased systemic blood pressure, or both. After a ∼1min imaging delay from administering C21, we observed a small increase in the HbO signal, followed by a slow decay back to baseline (Supplementary Figure 2B). Remarkably, spontaneous and stimulus-evoked hemodynamic responses were highly altered in Gq-DREADD mice in the presence of C21. In C21, there was a noticeable decrease in the number of spatiotemporally coordinated hemodynamic changes during null (no whisker stim) trials; instead, this was replaced by low-magnitude, window-wide, periodic fluctuations in signal intensity that repeated once every ∼8 seconds, demonstrating enhanced vasomotion by systemic C21 (Supplementary Figure 2C-D). Remarkably, the two-peak functional hyperemia response was largely abolished during whisker stimulation (Movie 4 and Movie 5); although a smaller change from baseline was still visible during S_1_, hemodynamic activity in S_2_ was severely curtailed and replaced instead by the same window-wide fluctuations in signal intensity seen in null trials (but of slightly higher magnitude). This effect was much more pronounced in HbR time-stacks than in HbO time-stacks. Inspection of behavior recordings showed that locomotion activity markedly reduced following C21 administration. However, the large C21 effect on hemodynamics could not be explained by this reduced running, since such a behavioral state still typically shows robust functional hyperemia (see Figure 3E). Furthermore, the mice still spontaneously sniffed, whisked, and groomed themselves during the whisker-puff trials, indicating the animals were not compromised.

**Figure 5:**
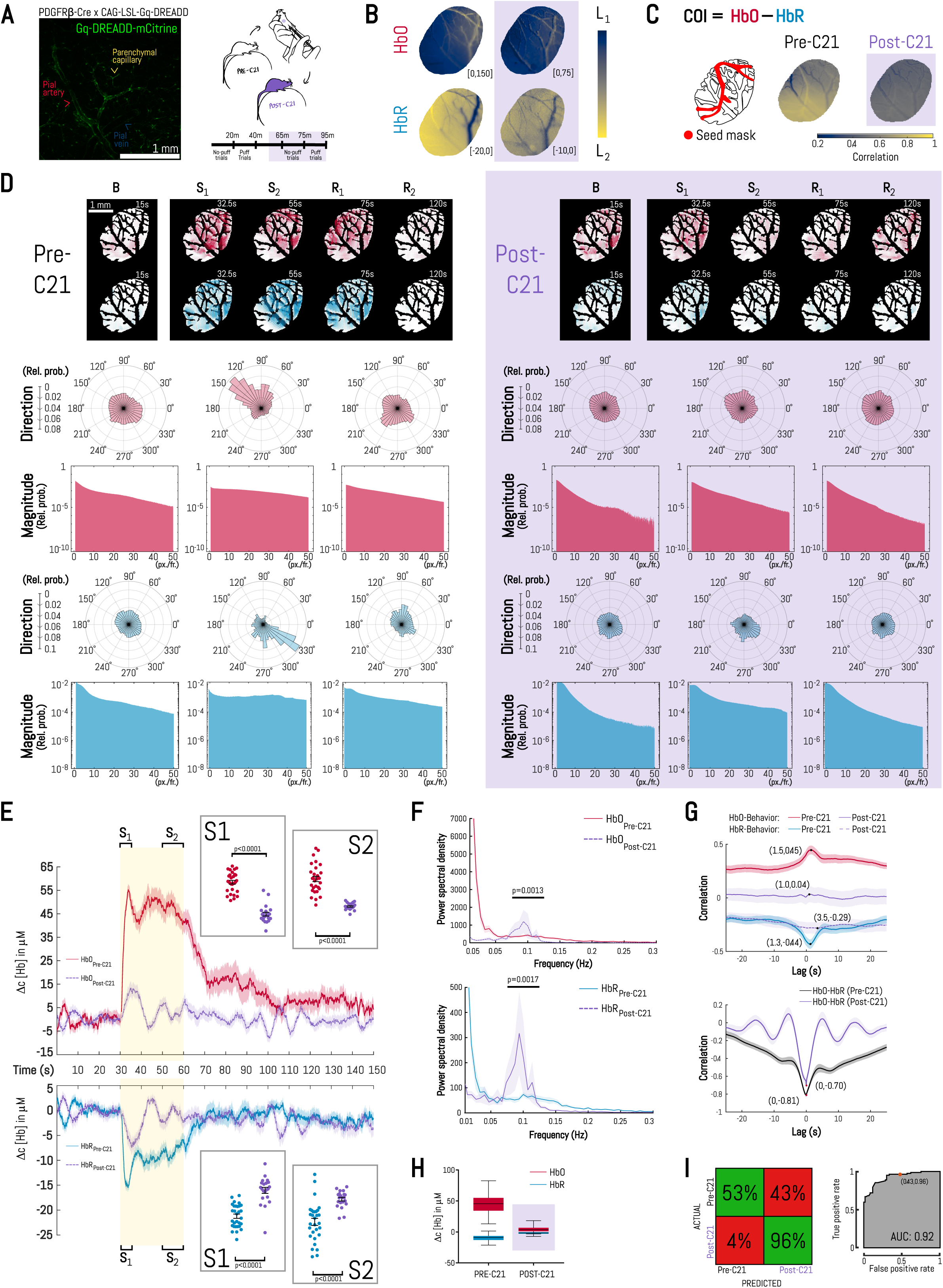
Direct vascular manipulation dysregulates functional hyperemia spatially and temporally. A. (Left) Barrel cortex from a PDGFRβ-Cre x CAG-LSL-GqDREADD-mCitrine mouse showing selective expression in mural cells (vascular smooth muscle cells and contractile pericytes); (right) Schematic of experimental protocol and timeline. Purple shading indicates the post-C21 period. B. Mean intensity projections of trial-averaged HbO (above) and HbR (below) time-stacks during whisker stimulation period (30-60 s) before and after systemic administration of 0.1 mg/kg Compound 21. Each image is an average of 5+ trials from the same animal. Colormap limits are provided in the bottom corner of each panel. Purple shading indicates post-C21 period. C. Results of seed-based correlation analysis on a representative trial-averaged COI time-stack before and after systemic administration of 0.1 mg/kg Compound 21: (left) seed-region; (right) correlation maps before and after drug administration. Colormap encodes correlation range [0.2, 1]. COI: Cerebral oxygenation index = HbO – HbR. Purple shading indicates the post-C21 period. D. Standard and polar histograms summarizing, respectively, magnitude and direction of optical flow vectors for 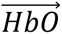 and 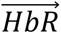 from trial-averaged time-stacks before (left) and after (right) administration of 0.1 mg/kg Compound 21. Purple shading indicates the post-C21 period. E. Trial-averaged time-courses of hemodynamic changes for puff trials before and after systemic administration of 0.1 mg/kg Compound 21. Data shown represents mean ± SEM. Yellow shading indicates whisker stimulus period. (Insets) Scatter dot plots showing distribution of trial means for HbO and HbR changes before (N=30 trials, 3 mice) and after (N=21 trials, 3 mice) systemic administration of 0.1 mg/kg Compound 21. Error bars: mean change ± SEM. Mann-Whitney two-tailed unpaired tests, HbO S_1_: U=12, P<0.0001; HbR S_1_: U=49, P<0.0001. Welch-corrected two-tailed unpaired t-tests, HbO S_2_: t(34)=10.09, P<0.0001; HbR S_1_: t(38)=6.98, P<0.0001; HbR S_2_: t(42)=5.66, P<0.0001. F. Time-courses of hemodynamic activity (left), and trial-averaged single-sided amplitude spectra of hemodynamic changes (right), for puff trials before and after systemic administration of 0.1 mg/kg Compound 21. Data shown represents mean ± SEM. Mann-Whitney two-tailed unpaired tests, HbO: U=150, P=0.0013; HbR: U=154, P=0.0017. G. (Left) Summary cross-correlation between hemodynamics and locomotion for puff trials before (above) and after (below) systemic administration of 0.1 mg/kg C21; (Right) Summary cross-correlation between HbO and HbR for puff trials before (above) and after (below) C21. Data shown is mean ± SEM. Yellow and purple shading indicate whisker stimulus, and post-C21 periods, respectively. H. Results of 2-way ANOVA between hemodynamic measure (HbO, HbR) and drug condition (Pre-C21, Post-C21). Interaction was significant (P<0.0001), accounting for 23% of the observed variance. Purple shading indicates the post-C21 period. I. (Above) Distribution of classification accuracies across 5771 features; (inset) confusion matrix for the linear SVM classifier used to separate hemodynamic time-courses from before and after Compound 21 administration. (Below) Low-dimensional representation showing the first two principal components of the classifier.

Inspection of mean intensity image projections of whisker-puff trials before and after systemic C21 reflected the increase in pial arteriole tone, and there was also a drastic reduction in the magnitude and spatial extent of hemodynamic activation within the cranial window (Figure 5B, note the different colormap limits). We performed seed-region correlation on trial-averaged time-stacks of the cerebral oxygenation index to visualize the connectivity between hemodynamic changes within pial arterioles and those elsewhere in the cranial window (Figure 5C). Comparison of these regional correlation maps before and after C21 administration showed a reduction in functional connectivity of feeding vessels with their tissue targets in all three mice, reflecting an uncoupling of oxygen supply to brain tissue. Finally, we performed optical flow analysis to better describe the observed changes in brightness patterns before and after C21 administration (Figure 5D). Following C21, flow patterns during the stimulation period became more turbulent, characterized by velocity vectors that were more uniform in size across the window and more prone to rapidly changing direction. Also apparent was the lack of a concentric response profile within the window in response to whisker stimulation following the vascular chemogenetic manipulation, which was more noticeable in S_2_ than in S_1_, and in HbR than in HbO time-stacks. Summary histograms reflect this striking disruption of stereotyped flow patterns: in C21, vector magnitudes appeared similar in baseline, stimulation and recovery periods, and there was a clear loss of directional bias.

We next performed detailed time and frequency domain analyses to further characterize this disruption of two-component functional hyperemia. Systemic administration of 0.1 mg/kg C21 significantly attenuated the stimulus-evoked response during S_1_ (p < 0.0001) and eliminated the clear distinction between S_1_ and S_2_ in a dose-dependent manner (Figure 5E, Supplementary Figure 3A); instead, we observed large, synchronized oscillations within the stimulation period in HbO and HbR time-series. These effects were absent in saline controls (Supplementary Figure 3B). Notably, the kinetics and single-sided amplitude spectra of the two-component response in naïve PDGFRβ × Gq-DREADD mice were dissimilar to that in PDGFRβ × GCaMP6s^+/–^ and PDGFRβ × GCaMP6s^−/−^ mice, which suggests that constitutive expression of the modified human muscarinic-3 receptor may affect intrinsic mural cell activity (Supplementary Figure 3C-D). Consistent with the data above, frequency domain analysis showed pronounced increases in spectral content within the ultra-low-frequency band (0.075-0.125 Hz) after C21 administration for both HbO (p = 0.0013) and HbR (p = 0.0017), and a suppression of all frequencies outside that window (Figure 5F). Cross-correlation analysis showed a decoupling of locomotion and whisking activity from hemodynamic activity following systemic C21 administration, yet the correlation between HbO and HbR still remained strongly negative (Figure 5G). Interestingly, two-way ANOVAs showed significant interaction between the hemodynamic measure (HbO, HbR) and drug groups (pre-C21, post-C21), showing a differential effect of C21 on HbO as compared to HbR (Figure 5H, p < 0.0001). Finally, to better understand the structure of pre-C21 and post-C21 time-series, we trained an unbiased, linear classifier to separate pooled HbO and HbR traces with >97% accuracy. The most discriminative features were measures of ‘stationarity’ (i.e., time-invariance of the time-series’ statistical properties) and ‘forecastability’ (i.e. ability to fit models to time-series data), followed by measures of ‘entropy’ and ‘spectral power’. Interestingly, the highest-performing features were related to the stability and predictability of the time-series: mean, variance and distribution within subsegments of the time-series; ‘one-step-ahead surprise metrics’; and measures of ‘multiscale entropy’ (i.e., dynamic complexity over multiple time scales).

## DISCUSSION

### Hemodynamic response to whisker stimulation in awake animals is a non-linear superposition of signals

Our data supports the idea that the hemodynamic response to a prolonged sensory stimulus in awake animals is a superposition of at least four signals: 1) infra-low-frequency (< 0.1 Hz) changes in vessel tone – likely related to changes in local metabolites or systemic factors such as blood pressure – which set the dynamic range of the tissue oxygenation response in response to local neuronal activity; 2) ultra-low-frequency (∼ 0.1 Hz) oscillations that are prevalent in the absence of a whisker puff stimulus; 3) behavior-related changes that are likely driven by the animal’s respiration cycle (∼ 2.5 Hz) and by neural activity related to natural whisking (Zhang et al. 2019); and, 4) a sustained, whisker stimulation-evoked change that exhibits a two-peaked increase in amplitude with stimulus duration and returns to baseline only ∼30 s after stimulus offset. We confirm previous reports of interaction between ongoing and stimulus-evoked cortical hemodynamics (Arieli et al. 1996, Fox et al. 2006, Saka et al. 2010), and propose that the interaction becomes distinctly non-linear under prolonged (>5s) sensory stimulation.

This nonlinearity arises from a complex interplay of local and systemic influences on brain microvessels that act in a spatially and temporally varying manner. We noticed that the kinetics of the evoked response was influenced significantly by the initial state properties of pial vessels; for example, the phase angle of oscillation immediately before stimulus onset affected S_1_ and S_2_ response magnitudes for HbO but only the S_2_ response magnitude for HbR. We noticed that slow-wave frequencies were amplified by whisker stimulation in a time-and frequency-dependent manner for HbO, but not for HbR. We also observed enhanced coupling between HbO (an oxygen ‘supply’ signal) and HbR (an oxygen ‘demand’ signal) during S_1_ and immediately after stimulus offset, which is likely related to the OFF response of thalamic barreloid neurons (Kyriazi et al. 1994). Such a spike in negative correlation between these signals suggests a transient increase in oxygen delivery (HbO – HbR), which reflects further supply-demand optimization during these periods. We observed that the duration of locomotion during a trial preferentially affected the magnitude of the late component of the hemodynamic response for HbO and HbR and shaped the frequency response in an ultra-low frequency window (0.075-0.125 Hz) for HbR but not for HbO. We also noted that overall variability in HbO and HbR activity was lower in the late recovery period (R_2_) than during the baseline period (B), even though the prevalence of spontaneous behaviors that contribute to the hemodynamic changes (natural whisking and locomotion) were no less prevalent during R_2_ than during B. This observation is consistent with the idea that prolonged sensory stimulation induces vascular ‘refractory periods’ in pial vessels (Cannestra et al. 1998). These periods of reduced vascular responsiveness also contribute to the proposed non-linearity because they 1) can vary in length, 2) can curtail the magnitude of subsequent stimulus-evoked and spontaneous responses, and 3) can be overcome by significantly novel or temporally spaced stimuli.

Importantly, and in line with a previous finding in cat visual thalamus (Rivadulla et al. 2011), we also showed that destabilizing the nonlinear dynamic equilibrium of ongoing hemodynamics can dramatically affect the spatial and temporal properties of neurovascular coupling in the cortex. Chemogenetic activation of mural cell Gq signaling via systemic C21 administration increased basal tone and intensified vasomotion. The effect on functional hyperemia was likely due to vasomotion, rather than pressure and/or vasoconstriction because 1) increases in blood pressure or arteriole tone, when they are acute, do not impair functional hyperemia (Blanco et al. 2008, Kazama et al. 2003, Iddings et al. 2015) and a previous *in vivo* study noted a reciprocal relationship between the prominence of vasomotion and functional hyperemia (Rivadulla et al. 2011). In C21, the magnitudes of HbO and HbR responses were severely curtailed, which would reduce total blood flow (HBO + HbR) and overall oxygen delivery (HbO – HbR). Furthermore, the onset of functional hyperemia appeared to synchronize vasomotion in C21 across the barrel cortex in all three animals because vasomotion-like rhythms ‘appeared’ in the averaged timeseries data; yet functional hyperemia could not reach its full amplitude or duration because it was quickly deflected into an oscillation. This dramatically decreased the spatial extent of activation, altered the functional connectivity of tissue oxygenation within the window, and virtually abolished the coordinated spatial response at the regional scale.

### Integrated spatial, temporal, and behavioral analyses are necessary to delineate heterogenous hemodynamic responses

Although WF-IOS imaging provides a wealth of two-dimensional hemodynamic data across time, few studies have systematically investigated the spatiotemporal properties of neurovascular coupling in an integrated way. We observed that slow, global changes in total blood flow during the baseline period in visually identified pial vessels sets the dynamic range of stimulation-evoked HbO and HbR responses in non-vessel regions of the cranial window. A negative-going HbT slope (which corresponds to a slow constriction) preferentially increased the magnitude of the late component (S_2_) with no effect on the early component (S_1_). This is consistent with a finding by Blanco and colleagues in rat cortical slices that the magnitude of the evoked diameter changes in pial arterioles is directly related to resting vascular tone (Blanco et al. 2008). A similar observation was made when HbO and HbR responses were categorized by pre-stimulus phase angle, which aligns with a previous finding by Saka and colleagues (Saka et al. 2012). Thus, cellular mechanisms responsible for the late component of functional hyperemia are strongly and preferentially influenced by the initial state of the vessel. This may further indicate that the same pathways involved in arteriole tone control are subsequently targeted during the late component of functional hyperemia.

We noted that 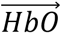 and 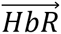 vector fields were opposite in sign, which was evident in vector field visualizations for individual trials, as well as in the summary polar histograms. Why would hemodynamics spread across the barrel cortex in this way? Indeed, if the inflow of oxygenated blood during functional hyperemia was ‘washing out’ deoxygenated hemoglobin molecules from areas of high metabolic activity, one should expect that the two signals would propagate in the *same* direction. Although recent reports imply that oxygen ‘leaks out’ of the walls of penetrating arterioles and flows down its concentration gradient in the tissue towards venules (Sakadzic et al. 2014), the random distribution of arterioles and venules in the barrel cortex should occlude such ‘oxygen countercurrents’ at the broad spatial scale of WF-IOS imaging. The most parsimonious explanation for this is that venular drainage may physically be running in a direction opposite to arteriolar supply. Indeed, feeding arteries enter anterolaterally into the barrel cortex, whereas feeding venules enter from the posteromedial aspect (Hartmann et al. 2018, Ma et al. 2016, Tran and Gordon 2015a, Xiong et al. 2017). Although the literature concerning the relative orientation of arterioles and venules and the resultant distribution patterns of blood flow is rather limited, recently published cerebral vascular maps of the mouse brain (Blinder et al. 2013, Xiong et al. 2017) suggest that angio-architectural features of the neocortex overlaid arterial and venous anastomoses on the cortical surface, penetrating vessels that run in parallel, and a tortuous mesh of subsurface microvessels, can at a network level produce antiparallel flow patterns.

We noted that the distinction between the hemodynamic response in center (active) and surround (passive) regions was stronger for HbO than for HbR, and it was more pronounced during S_1_ than during S_2_. Moreover, we noted that locomotion and associated natural whisking can modulate (but does not dominate) this center-surround effect in a spatially non-specific manner. Additional studies of neurovascular coupling that employ such integrated spatiotemporal approaches in awake animals are needed to improve our understanding of this complex and essential phenomenon.

### Separate control mechanisms for cortical blood flow during prolonged sensory stimulation

We speculate that separate cellular mechanisms operating on distinct timescales and exhibiting differential regulation likely drive the two-peak oxygenation response. For example, the activity of neurons embedded in capillary beds drives the initial increase in blood flow (Lee et al. 2010); thus, cell pathways in excitatory and/or different classes of inhibitory neurons are key to at least the early component of the response (Echagarruga et al. 2020, Lee et al. 2020). There is ample evidence that astrocytes exhibit a delayed (∼3-5 s) elevation in free Ca^2+^ after the onset of neural activity (Ding et al. 2013, Tran et al. 2018, Stobart et al. 2018, Gee et al. 2014, Wang et al. 2006, Lind et al. 2013, Nizar et al. 2013), which can be potentiated by movement and modulated by norepinephrine (Ding et al. 2013, Paukert et al. 2014, Polack et al. 2013, Tran et al. 2018). Thus, astrocytes may participate in the later component of the blood flow response as they are influenced by locomotion state and because astrocyte Ca^2+^ signals have been correlated with the late, non-linear component of BOLD fMRI responses to sustained brain activation (Schulz et al. 2012). Notably, under anesthesia, astrocyte Ca^2+^ transients are suppressed (Thrane et al. 2012) and arteriole/CBF responses to sustained sensory stimulation consist of an initial rise followed by a plateau at close to the same amplitude (Lacroix et al. 2015, Lecrux et al. 2011, Liu et al. 2008, Peng et al. 2002, Toth et al. 2015). But here, we see a clear two-component response in awake animals. Finally, the microvasculature itself has distinct temporal components: an early vasodilation from local dilators acting on mural cells and fast-acting electrical conduction along the endothelium (Chaigneau et al. 2003, Chen et al. 2014, Hall et al. 2014, Longden et al. 2017), followed by a slower conducting wave of Ca^2+^ and nitric oxide along the endothelium (Budel et al. 2003, Tallini et al. 2007). These separate cell-types and cellular mechanisms operating on distinct timescales likely underlie the spatiotemporal cerebral hemodynamics we observed to prolonged whisker stimulation.

### Implications for functional imaging

Functional magnetic resonance imaging (fMRI) studies routinely employ paradigms involving prolonged sensory stimulation, and typically interpret BOLD changes as convolutions of a linear kernel with underlying neural activity. This is problematic because during sustained neural activation, the BOLD response becomes nonlinear: more, or less, oxygen is delivered locally than should be reasonably expected given the measured neural activity. Here, we provided evidence that prolonged sensory stimulation evokes a complex blood flow response, the later component of which is more apt to dynamic regulation by the initial conditions of the vasculature. We also showed that behavioral state can affect the magnitude, frequency, and spatial profile of the stimulus-evoked response, which may be a confound of vascular origin for inferring neural activity from hemodynamic signals. We thus reaffirm the need for developing more nuanced transfer functions linking BOLD activity to underlying neural activity.

Unlike fMRI, functional near-infrared spectroscopy (fNIRS) can resolve changes in HbO and HbR. These two hemodynamic measures are assumed to be negatively correlated, and the cross-correlation of the HbO and HbR signals has been recently proposed as a feature to characterize dynamic noise and reject motion artifacts in fNIRS imaging (Lee et al. 2018). Yet, we provided evidence here that the magnitude, frequency, and spatial extent of HbO and HbR responses can be independently modulated, which demands a more careful assessment of that assumption, especially in low signal-to-noise conditions.

### Limitations of the study

Bone and dura were removed during surgical preparation, which eliminated signal contributions from extracortical vessels and improved the penetrance of illumination light into the cortex. However, this can alter intracranial pressure and/or perturb constitutive dura-mediated signaling (Mastorakos and McGavern 2019), both of which may conceivably affect NVC as a result of altered mechanical equilibrium within the cortex (Gao et al. 2015) and/or via systemic signaling pathways. Moreover, imaging with a glass window or even a thinned brain surface may cause cooling of brain tissue, which can affect several factors related to blood flow and tissue oxygenation (e.g. vasoconstriction, O_2_ diffusion and solubility, reduced hemoglobin affinity to O2) (Roche et al. 2019).

There was a slight motion artifact at the onset of whisker stimulation and at the onset of locomotor activity. We did not employ image registration to correct for such artifacts; instead, data from the first 500 milliseconds of whisker stimulation was discarded. Nonetheless, this period should be similar to baseline data as functional hyperemia takes ∼700ms to initiate.

WF-IOS imaging reports a superficially weighted sum of intrinsic signals from a volume of brain tissue. Furthermore, when measuring over the entire barrel cortex, hemodynamic changes in large feeding/draining vessels contribute to the measured changes in HbO and HbR. Since retrograde propagation of endothelial hyperpolarization proceeds at ∼2mm/s and causes smooth muscle relaxation in upstream branches (Chen et al. 2011), the contribution of feeding vessels likely increases over the course of the 30-second stimulation. Thus, the observed signal is a complex mixture of vascular activity in different compartments and at different depths of cortex and the phenomena reported (e.g. behavioral-state-dependence of tissue oxygenation) may vary by vascular compartment and/or tissue depth.

We quantified locomotion using an image-difference approach of the ball treadmill surface frame to frame. Although the temporal resolution (50 Hz) rendered it effective for aligning hemodynamic signals to running onset and for cross-correlation analyses, this measure of mouse locomotion is relative and non-specific.

## METHODS

### Animals

All animal procedures were approved by the University of Calgary’s Animal Care and Use Committee (AC19-0170). All studies were performed on male, P35-60, Pdgfrβ-Cre × RCL-GCaMP6s mice (n=21) or Pdgfrβ-Cre × LSL-CAG-Gq-DREADD mice (n=4) weighing between 21-30 g. Mice were kept on a 12hr:12hr light-dark cycle (ZT0 = 6AM) with food and water available *ad libitum,* and were group-housed until head-bar installation at P30-45. Five mice were excluded from detailed spatiotemporal analyses owing to visible damage from surgery and/or obvious movement artifacts due to imperfect cranial window installation.

### Surgeries

All surgeries used standard aseptic procedures and isofluorane anesthesia (5% for induction and 1- 1.5% for maintenance of reflex suppression during surgery; Pharmaceutical Partners of Canada Inc., Richmond Hill, ON, Canada). During surgery, mice were mounted in a stereotaxic frame and body temperature was maintained at 37°C using a heating blanket with a feedback rectal probe (Homeothermic blanket monitoring system, Harvard Apparatus). A single dose of buprenorphine (0.05 mg/kg, subcutaneous; controlled drug supplied by the University of Calgary’s Animal Resource Centre) was given at the start of each surgery. <u>Headbar-installation</u> One week before imaging, a minor surgery was performed to install a custom-made, stainless steel headbar. Briefly: the fur, skin and periosteum covering both hemispheres of the skull were removed; the headbar was positioned over the interparietal bone and glued down with cyanoacrylate glue (3M Vetbond™, 3M Animal Care Products, St. Paul, MN, USA) and dental cement (Ortho Jet Powder and Liquid, Lang Dental Manufacturing Co., Wheeling IL USA). A single dose of enrofloxacin (2.5 mg/kg, subcutaneous; Bayer Health Care, Toronto, ON, Canada) was given at the end of surgery to prevent post-operative infection. Mice were single housed in the days leading up to imaging to allow for safe recovery and for headbar stability on imaging day. <u>Craniotomy</u> This procedure was adapted from previously published protocols (Holtmaat et al. 2009, Mostany and Portera-Cailliau 2008, Muniak et al. 2012, Shih et al. 2012), and is described in full in a previous publication (Tran and Gordon 2015a). Briefly: a 3×3mm circular window was drawn over the whisker barrel cortex with a fine tip marker; the area of skull immediately adjacent to the circle was thinned down until a bone island forms; a fine tool was then used to lift the entire bone island off the skull; a small incision was then made in the dura mater using a 30 gauge needle, taking care not to touch the cortical surface, and the dura mater over the whisker barrel cortex was carefully removed; a pre-cut piece of cover glass was placed over the exposed cortex, and then glued down with cyanoacrylate glue at the edges. Drilling, dura removal and coverslip placement were all done under artificial cerebrospinal fluid to keep the brain surface cool and wet, and care was taken not to prevent the glue from coming into contact with the exposed cortex.

### Awake-mouse imaging

The air-supported spherical treadmill, which is critical for fully-awake and head-restrained imaging, was adapted from a previously published protocol (Dombeck et al. 2007) and a detailed description is given in a previous publication (Tran and Gordon 2015a). During habituation and imaging, mice received a 30-second air puff to contralateral vibrissae once every ∼5 minutes from the Picospritzer III and were exposed to the 10 Hz strobing illumination of red and green LEDs. During the inter-trial interval (∼2 minutes), illumination and air puff stimuli were absent. <u>Training</u> To reduce the stress associated with prolonged head-fixation and whisker stimulation, mice received 2-3 training sessions on the experimental apparatus in the days leading up to acute imaging. Once head-fixed, mice immediately began running on the spherical treadmill; this is initially erratic but becomes more controlled with experience. Training sessions were ∼75 minutes long, and the conditions were identical to those during acute imaging. They consisted of an initial 15-minute familiarization period (no air puff), followed by 60 minutes of whisker stimulation sessions. <u>Acute imaging</u> Following the craniotomy, the mouse was immediately transferred to the spherical treadmill and was allowed to recover for ∼45 minutes. Acute imaging sessions lasted between 60 and 90 minutes.

### Pharmacology

For experiments involving the user of designer receptors exclusively activated by designer drugs (DREADDs), the subject first underwent a complete imaging session (5 null trials and 5 or more whisker-puff trials). The mouse was then quickly removed from the ball and given an intraperitoneal injection (<0.1 mg/kg) of the water-soluble DREADD agonist Compound 21 (C21, Hello Bio: HB6124). The mouse was then returned to the ball for another imaging session (drug arrival, 5 null trials, 5+ whisker-puff trials). The experimenter was not blinded to the treatment. Since the effects of this manipulation were very pronounced, only three animals per group were required to reach statistical significance; thus, all statistical analyses were performed using pooled trials in each group.

### Instrumentation

We custom-built a widefield intrinsic optical signal imaging setup consisting of a sensitive camera (Basler Ace U acA1920-40 µm with Fujinon: HF50XA-5M lens; QE = 70%, well depth ∼33 ke-, dark noise ∼7e-, SNR ∼45dB; frame rate = 40 Hz, pixel size = 5.86 µm), stable light-emitting diodes (M530L3 and M660L4, Thorlabs Inc.), and filtering (Semrock) and collimation optics (Thorlabs). The lens was mounted 1 inch away from the camera to get a reduced field-of-view at a similar working distance, and 4×4 binning was used to improve signal-to-noise; this led to an effective pixel size of 23.44 µm. For behavioral monitoring, we used near-infrared illumination (780 nm LEDs) and two infrared-sensitive cameras; Camera 1 (40 Hz) captured movements of a small region of the Styrofoam ball, and Camera 2 (50 Hz) captured mouse behaviors like running and grooming from the front. An open-source pulse train generator (Sanworks, Pulse Pal v2) was configured to synchronize trigger signals between the cameras, LEDs and the Picospritzer III.

### Data analysis

#### WF-IOS imaging

Reflected red and green light from the cortex was collected at a sampling rate of 10 Hz per channel. Raw data consisted of 2D red- and green-channel reflectance images from each recording session (Supplementary Figure 4). These 3D ‘time-stacks’ of reflectance were then converted into 3D oxyhemoglobin (HbO) and deoxyhemoglobin (HbR) time-stacks of concentration using a custom MATLAB script based on the relationships described by Dr. Elizabeth Hillman’s group (Ma et al. 2016). Time-stacks of total hemoglobin (HbT = HbO + HbR) and cerebral oxygenation index (COI = HbO -HbR) were created via pixel-by-pixel addition or subtraction of HbO and HbR time-stacks. For the modified Beer-Lambert relation, we used scattering-adjusted pathlengths of 4.535 mm and 0.411 mm for red and green light, respectively (Ma et al. 2016). HbO and HbR time-stacks were averaged along the two spatial dimensions to create 1D HbO and HbR ‘time-series’ by computing the mean gray value for each image. Unless otherwise specified, this averaging was performed for every pixel in the image. For whisker stimulation trials, time-stacks and time-series were normalized by subtracting baseline activity (0-30 s) on a pixel-by-pixel basis. Frames from the first 500 ms of the whisker stimulation period were excluded from all analyses due to the presence of obvious motion artifacts that caused an increase in red and green reflectance.

#### Time-series variability

Two different approaches were used to characterize the variation in HbO and HbR time-series without removing outliers. First, Poincaré return maps were created for each whisker stimulation trial by plotting every data point in the temporal segment of interest against the successive data point. For accurate comparison, we used segments of equal length (B: 0 – 30 s; R_2_: 105 s – 135 s). Maps for all trials were then overlaid on the same axis, unweighted least-squares fitting was performed for the conic representation of an ellipse with possible tilt (Fitzgibbon et al. 1996, Gal 2003), and standard estimates of dispersion (SD1, SD2 and 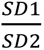) were computed for each temporal segment (Fishman et al. 2012). Second, we computed coefficients of variation 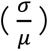 for each trial from the temporal segment of interest.

#### Frequency domain analysis

A discrete Fourier transform was computed for each hemodynamic time-series and the power spectral density (PSD) was computed in a broad range, [0.01 5] Hz. To visualize spectral changes as a function of time, spectrograms were created by computing Hamming-windowed short-time Fourier transforms for 6-second segments of each hemodynamic time-series with 95% overlap between segments. Individual PSD profiles and spectrograms were averaged on a trial-by-trial basis to facilitate comparisons between groups. To average trials by pre-stimulus phase-angle, we replicated the approach described by Dr. Jason Berwick’s lab, wherein the Hilbert transform is used to calculate the instantaneous phase of pre-stimulus fluctuations in hemodynamic activity (Saka et al. 2010). We defined the pre-stimulus period as being the 500 ms preceding whisker stimulus onset and grouped phase-angles as rising (-90° to 90°) or falling (90° to 270°).

#### Moving correlation

Pearson product-moment correlation coefficients were computed between HbO and HbR time-series with a 6-second sliding window, and missing values at endpoints were discarded. Since the correlation was persistently high and negative, heatmap color limits for the R-value were set to [-1 -0.75].

#### Curve-fitting

Time constants (τ) were calculated from first-order exponential fitting.

#### Image processing

HbO and HbR time-stacks were averaged along the time dimension to create 2D mean intensity projections; doing this for specific time windows (e.g. S_1_: 30-35 s) permitted visual comparison of spatial extents of hemodynamic response as well as the identification of large/medium-sized vessels within the cranial window. Binary masks were created by using the ‘Adjust Threshold’ function in ImageJ or by directly replacing values in the MATLAB image matrix with 0 or 1 (for exclusion or inclusion, respectively).

#### Seed-region correlation

Our implementation extends a seed-based correlation approach described in a publication from Dr. Timothy Murphy’s lab (Silasi et al. 2016). Briefly: all pixels within the initial seed mask are registered; pixelwise linear correlation is performed between each pixel in the mask and all other pixels within the window; the resulting correlation matrices (one for each pixel in the initial seed mask) are averaged to create the raw seed-region correlation matrix; all entries of the seed-region correlation matrix that fail to reach a histogram-based threshold (e.g. 75^th^ percentile) are set to zero.

#### Optical flow analysis

Velocity vector fields were estimated from individual or trial-averaged HbO and HbR time-stacks using MATLAB’s ‘opticalFlowFarneback’ function, which estimates the direction and speed of moving elements in video sequences using the Farnebäck method (Farnebäck 2003).

#### Locomotion analysis

To quantify locomotion activity from recordings of a small region of the Styrofoam ball, we implemented a simple algorithm in ImageJ (Supplementary Figure 5): briefly, the image stack was read in and duplicated; the first frame from ‘Stack1’ and the last frame from ‘Stack2’ were deleted; a pixel-by-pixel subtraction was performed for every image in the stack (i.e. Stack1 – Stack2); mean gray value was plotted against time for the resulting stack. Where required, the MIJ package (Sage 2012) was used to exchange images between MATLAB and ImageJ.

#### Machine learning

We used HCTSA (a MATLAB-based software package) to extract 5000+ normalized features from HbO and HbR time-series (Fulcher and Jones 2017, Fulcher et al. 2013). A Support Vector Machine (SVM) classifier (linear kernel; tenfold cross-validation) was subsequently trained using MATLAB to separate meaningful groups of time-series (‘Center’ vs ‘Surround’, ‘Before C21’ vs ‘After C21’). Classification performance was assessed using standard metrics (confusion matrix, receiver operating characteristic curve, permutation test relative to a shuffled ensemble, sensitivity to specific feature sets, and comparison to lower-dimensional feature spaces). We also used DeepLabCut v2.1.10.2 (Mathis et al. 2018, Nath et al. 2019, Mathis et al. 2019) to perform marker-less position-tracking of two or three visible whiskers in each frame. We used the single-animal implementation of the deep-learning approach with each whisker defined as a connected body with three parts (base, midpoint, tip). We manually labeled ∼120 images per animal sampled across a number of trials, and trained individual neural networks using GPU-based hardware acceleration provided by Google Colab with the following parameters: TrainingFraction=0.95; net_type=resnet_50; augmenter_type=imgaug; max_iters=200000. We assessed tracking performance by examining the labeled videos and the filtered x/y-position traces. For follow-up cross-correlation and frequency domain analyses, we generated whisking-activity traces either by averaging x- and y-position traces.

### Statistical analysis

All analyses were performed in GraphPad Prism 7.04. Checks of normality (D’Agostino-Pearson K^2^ test) and equal variance (F-test) were first performed on all data sets, and the appropriate two-tailed parametric or non-parametric test was selected. When comparing means from two matched groups, paired t-tests or Wilcoxon matched-pairs signed rank tests were used. When comparing means from two unmatched groups, unpaired t-tests (with Welch’s correction if variances were non-equal) or Mann-Whitney U-tests were used. When comparing means of multiple groups, Greenhouse-Geisser-corrected repeated-measures one-way analysis of variance (with Sidak’s correction for multiple comparisons) or Friedman tests (with Dunn’s correction for multiple comparisons) were used. All tests were conducted to α=0.05 significance level. No explicit blinding or randomization was performed. Summary data in-text are reported as mean ± standard error of the mean (SEM).

## RESOURCE AVAILABILITY

Further information and requests for resources and reagents should be directed to and will be fulfilled by the Lead Contact, Grant R Gordon.

## ACKNOWLEDGEMENTS

This work was supported by the University of Calgary, the Hotchkiss Brain Institute and Canadian Institutes of Health Research. G.R.G. was supported by Canada Research Chairs. We thank Rodney Barasi and Dr. Frank Visser for assistance with breeding and genotyping of transgenic mice. We thank Gayathri R Subramani for assistance with labeling images and training the deep learning model.

## AUTHOR CONTRIBUTIONS

Conceptualization, G.P. and G.R.G; Methodology, G.P., L.Y., K.M. and G.R.G; Software, G.P., L.Y. and K.M.; Formal Analysis, G.P.; Investigation, G.P.; Resources, G.P. and K.M.; Writing – Original Draft, G.P.; Writing – Review & Editing, G.P., K.M. and G.R.G.; Visualization, G.P.; Supervision, G.R.G.; Project Administration, G.R.G.; Funding Acquisition, G.R.G.

## DECLARATION OF INTERESTS

The authors declare no competing interests.

## SUPPLEMENTAL INFORMATION

**Movie 1**: Spatial dynamics of HbO changes (left) and HbR changes (right) during 30-second whisker stimulation for a representative trial (Mouse 7, Trial 6).

**Movie 2:** Spatial dynamics of the 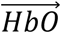 vector field during 30-second whisker stimulation for a representative trial (Mouse 7, Trial 6).

**Movie 3:** Spatial dynamics of the 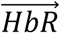 vector field during 30-second whisker stimulation for a representative trial (Mouse 7, Trial 6).

**Movie 4:** Spatial dynamics of HbO changes before (left) and after (right) systemic administration of 0.1 mg/kg Compound 21.

**Movie 5:** Spatial dynamics of HbR changes before (left) and after (right) systemic administration of 0.1 mg/kg Compound 21.

**Supplementary Figure 1:**
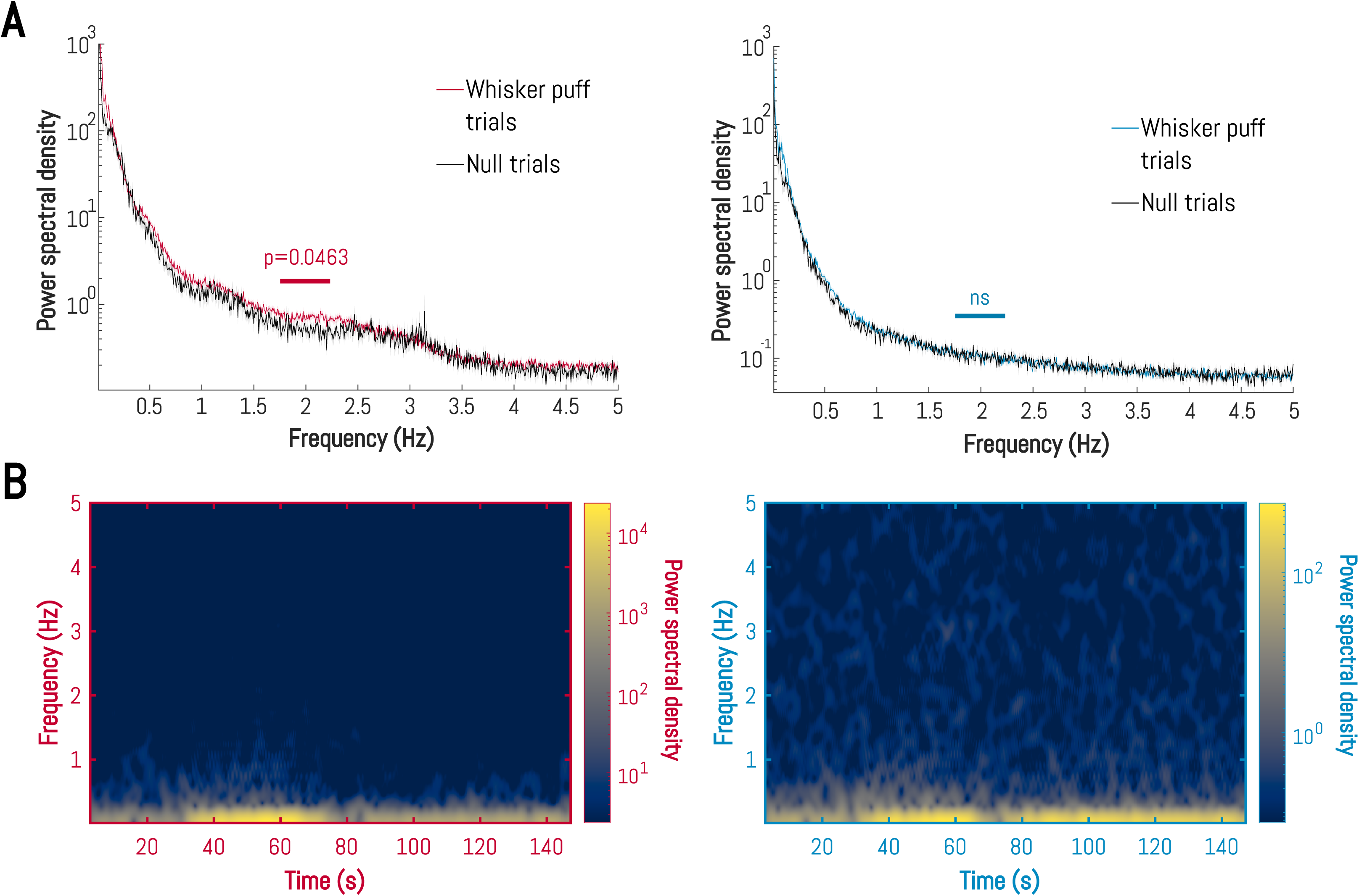
Detailed frequency domain comparisons of null and whisker puff trials. A. (Above) Trial-averaged single-sided amplitude spectra of HbO changes (left) and HbR changes (right) for null and whisker puff trials. Statistical comparison reported is made between the single-sided amplitude spectra of null trials and those of whisker puff trials in the frequency range [1.75,2.25] Hz. Paired two-tailed t-tests, HbO: t(14)=2.19, P=0.0463; HbR: t(14)=0.98, P=0.3434. ‘ns’: not significant B. (Above) Trial-averaged spectrograms of HbO (left) and HbR (right) activity during whisker puff trials.

**Supplementary Figure 2:**
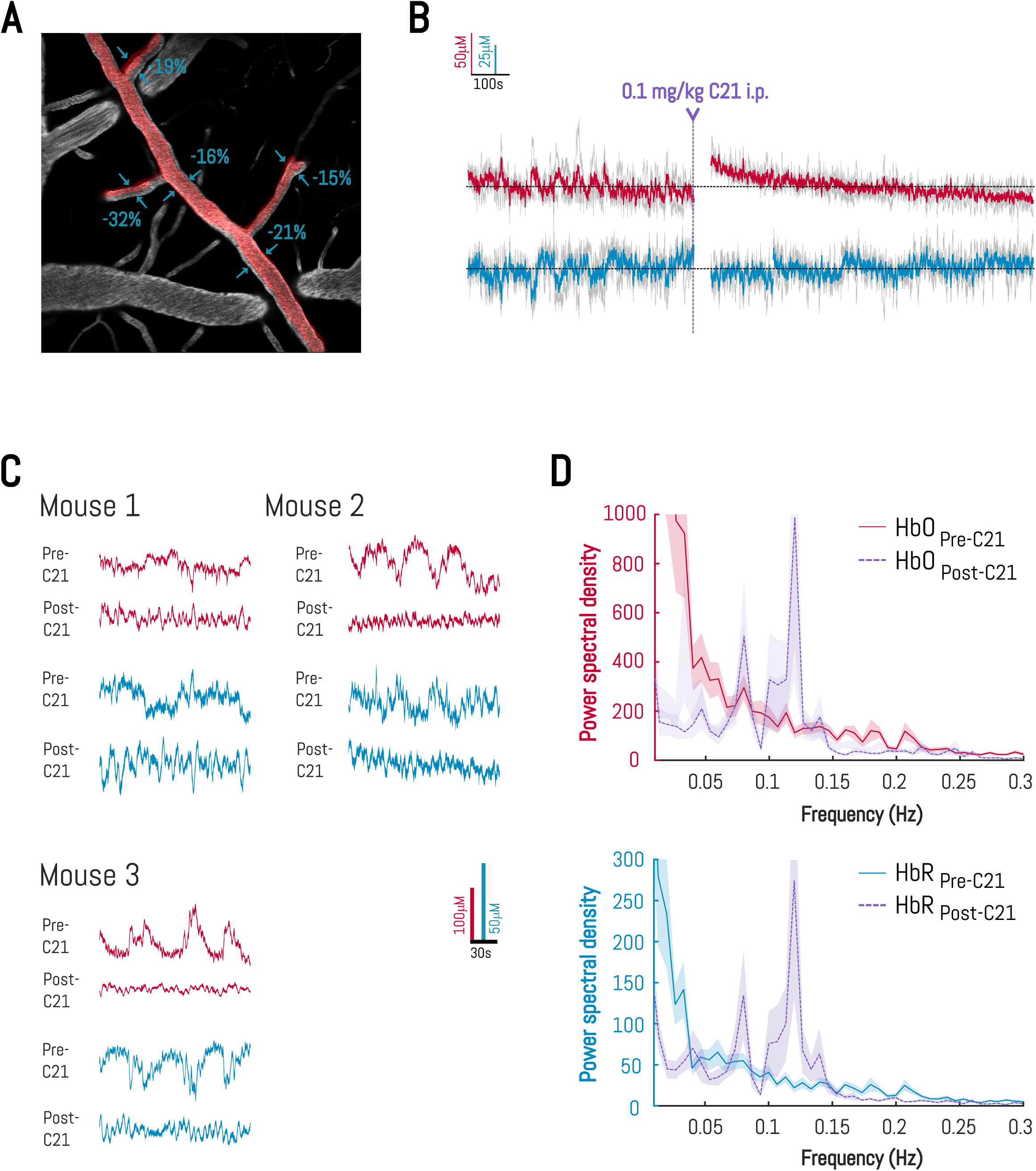
Change in basal tone and vasomotion following C21 administration. A. Response of a single pial arteriole to 0.1 mg/kg C21 measured by two-photon fluorescence imaging in an awake mouse. Red overlay of pial arteries shows increased arterial tone after 900 s, and blue text indicates percentage change in vessel diameter. B. Time-course of hemodynamic changes immediately before, and 45 seconds after, 0.1 mg/kg Compound 21 injection. Color trace is mean, gray traces are overlaid raw data from 3 animals. Purple dotted line indicates timepoint of C21 injection. ‘i.p.’: intraperitoneal C. Representative time-courses of hemodynamic changes for null trials from each animal before and after systemic administration of 0.1 mg/kg Compound 21. D. Trial-averaged single-sided amplitude spectra of hemodynamic changes for null trials. Data shown represents mean ± SEM.

**Supplementary Figure 3:**
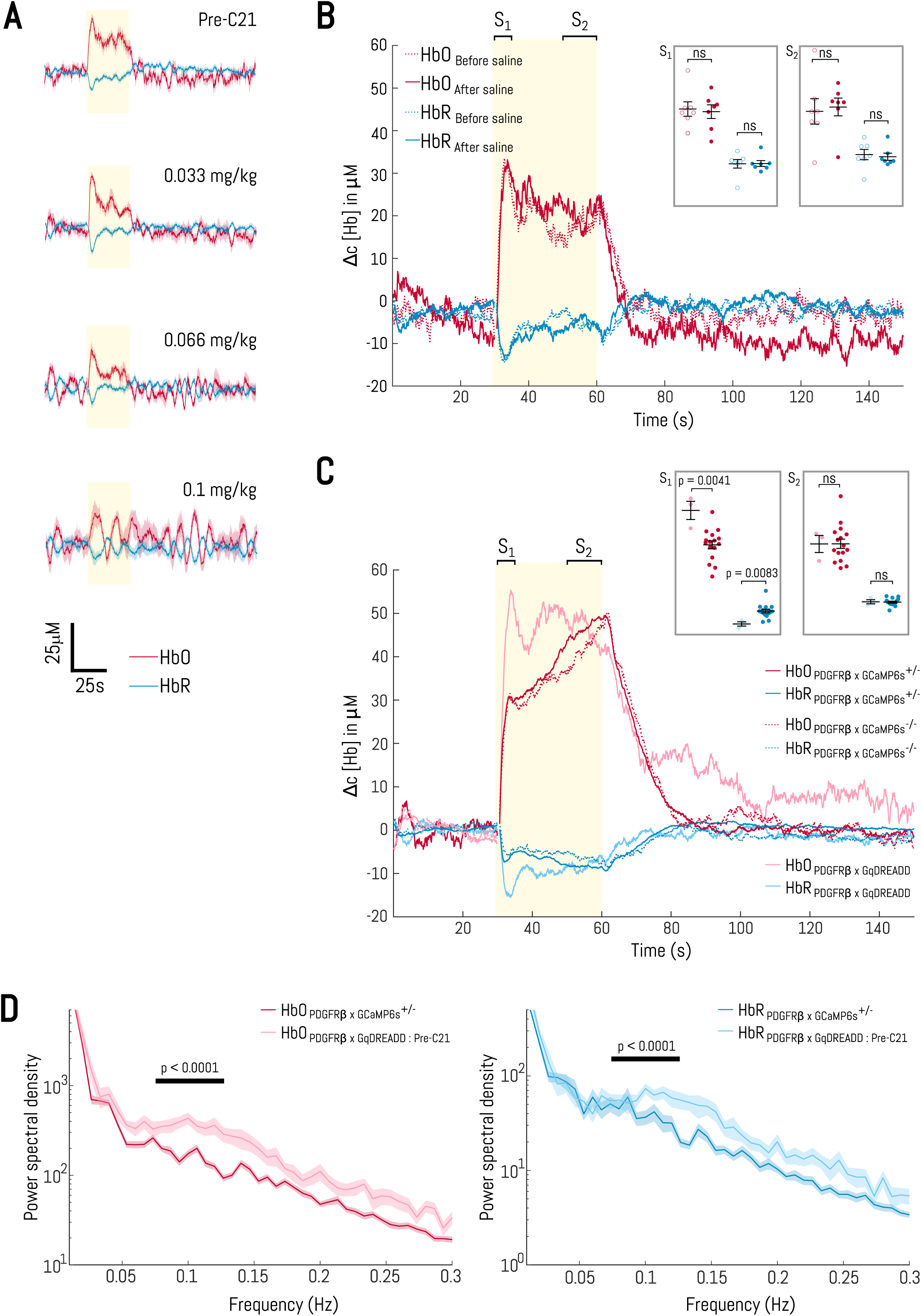
Control experiments for the GqDREADD manipulation. A. Trial-averaged time-courses of hemodynamic changes during whisker-puff trials in one mouse before and after systemic Compound 21 administration: *(Above)* 0.033 mg/kg; *(middle)* 0.066 mg/kg; *(below)* 0.1 mg/kg. Purple shading indicates post-C21 trials. B. Trial-averaged time-courses of hemodynamic changes for puff trials before and after systemic administration of saline. Data shown represents mean ± SEM. Yellow shading shows whisker stimulus period (Insets) Scatter dot plots showing distribution of trial means for HbO and HbR changes before (N=7 trials, 3 mice) and after (N=7 trials, 3 mice) systemic administration of saline. Error bars: mean change ± SEM. Wilcoxon two-tailed matched-pairs signed rank tests, HbO S_1_: W=4, P=0.8125; HbO S_2_: W=12, P=0.3750; HbR S_1_: W=-8, P=0.5781; HbR S_2_: W=-14, P=0.2969. C. Time-courses of hemodynamic changes in naïve Pdgfrβ x GqDREADD, Pdgfrβ x GCaMP6s^+/–^ and Pdgfrβ^+/–^ x GCaMP6s mice. Data shown are mean traces. (Insets) Scatter dot plots showing distribution of trial means for HbO and HbR changes in genotype-positive Pdgfrβ x GqDREADD (N=30 trials, 3 mice) and Pdgfrβ x GCaMP6s (N=214 trials, 16 mice) animals. Error bars: mean change ± SEM. Mann-Whitney two-tailed unpaired test, HbO S_1_: U=1, P=0.0041; HbO S_2_: U=22, P=0.8751; HbR S_1_: U=2, P=0.0083; HbR S_2_: U=22, P=0.8751. D. Trial-averaged single-sided amplitude spectra of hemodynamic changes in Pdgfrβ x GqDREADD mice, and in Pdgfrβ x GCaMP6s^+/–^ mice, before systemic Compound 21 administration. Mann-Whitney two-tailed unpaired test, HbO: U=3277, P<0.0001; HbR: U=0, P<0.0001

**Supplementary Figure 4:**
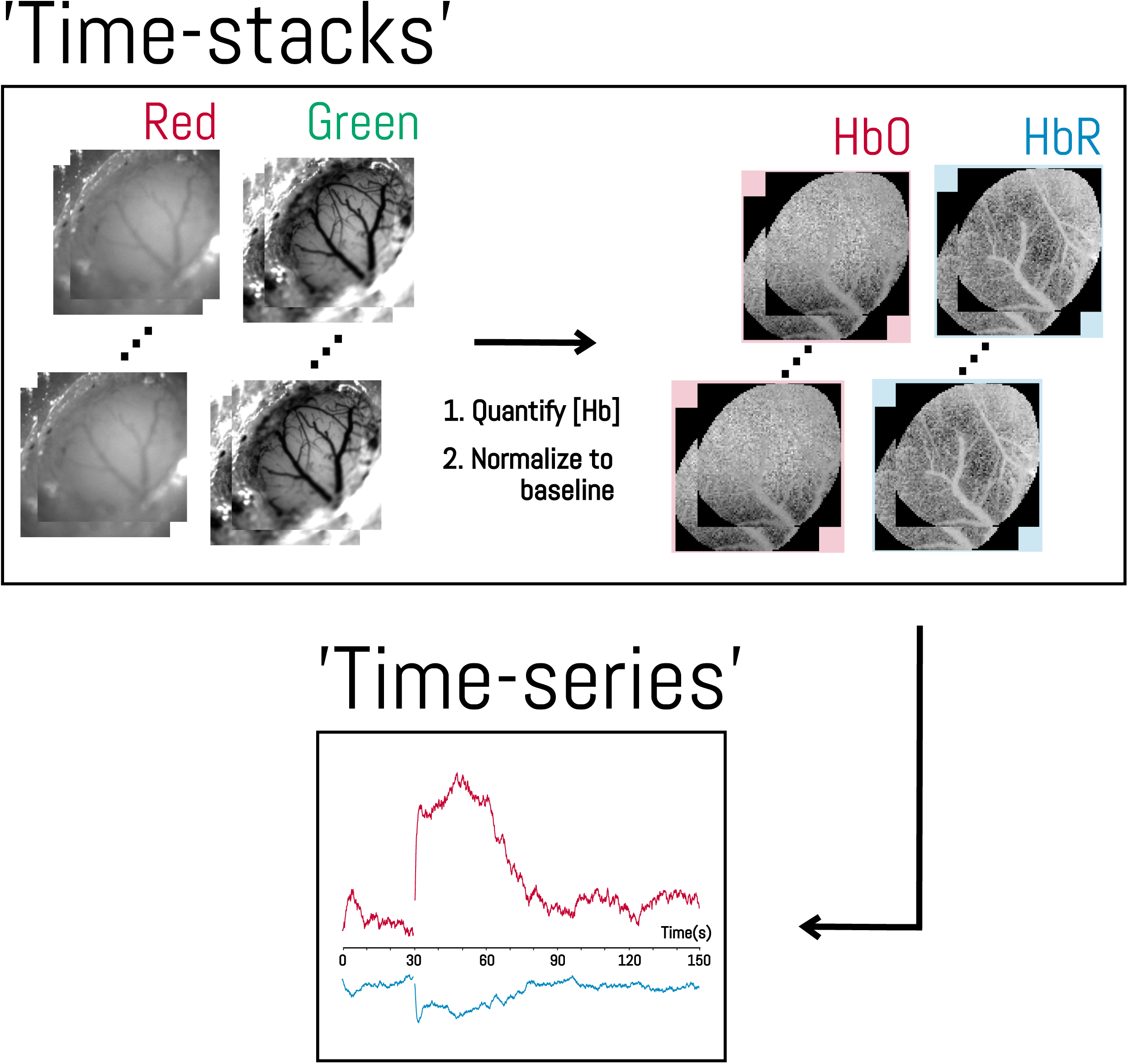
Time-stacks and time-series are used for spatial and temporal analyses, respectively.

**Supplementary Figure 5:**
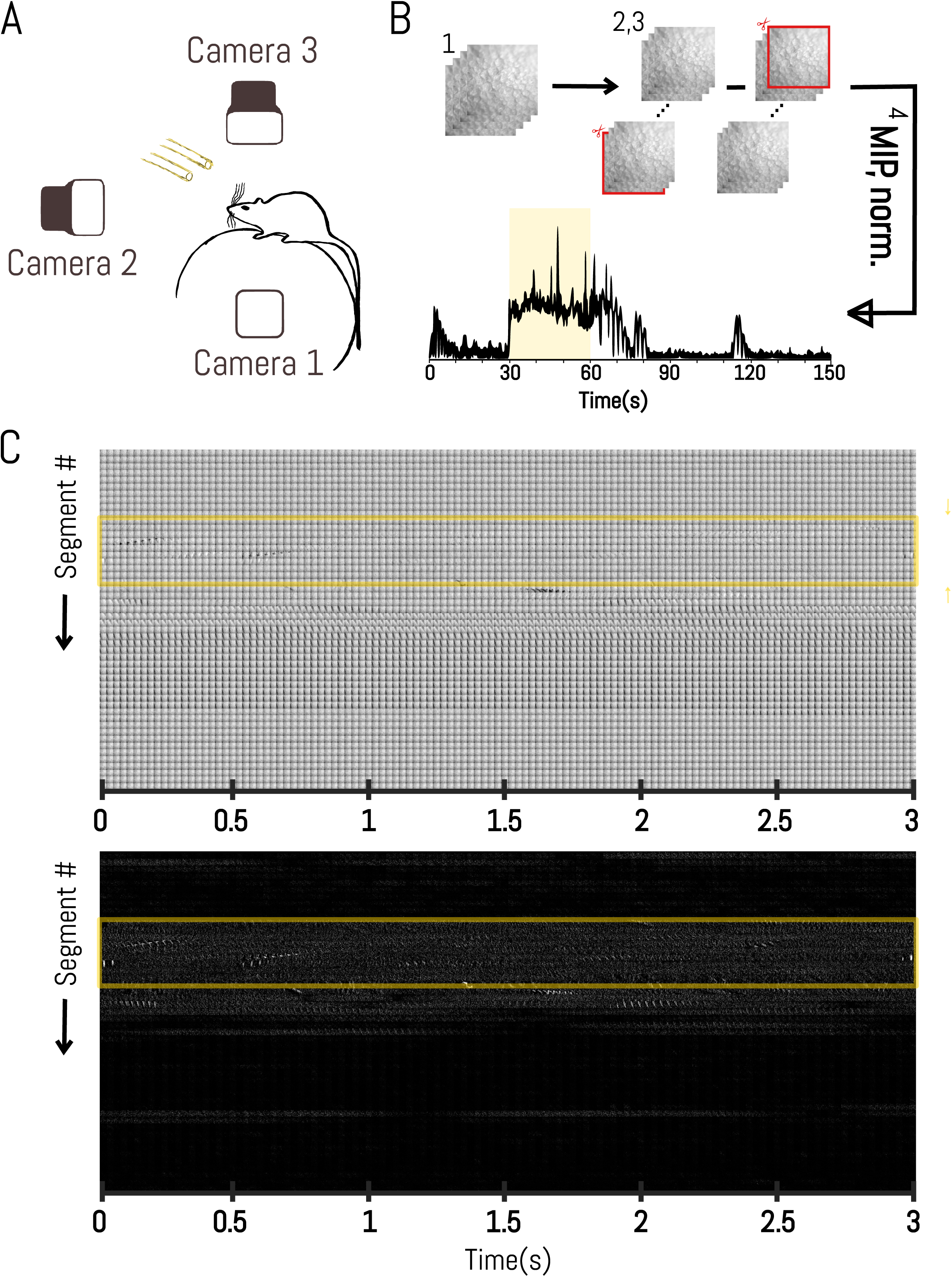
Mouse locomotion can be quantified from recorded treadmill movements using image differencing. A. Schematic of behavioral monitoring setup. B. Steps of difference-image analysis. Briefly: 1) Import image stack; 2) Duplicate image stack; 3) Delete first slice of first duplicate and last slice of second duplicate and subtract second stack from first; 4) Compute each frame’s mean gray value and normalize to baseline. ‘MIP’: mean intensity image projection; ‘norm.’: normalization. C. Montage of raw frames from Camera 1 (above) and difference-images (below) from a single puff trial (‘Mouse 10, Trial 6’). Each row represents a 3-second segment from the puff trial. Yellow box indicates whisker stimulation period.

## Notes

### Competing Interest Statement

The authors have declared no competing interest.

